# A sampling-theory framework for richness–uniqueness relationships

**DOI:** 10.64898/2026.10.04.756530

**Authors:** Xiaoning Wang, Dingliang Xing

## Abstract

Ecological uniqueness, commonly quantified by local contribution to beta diversity (LCBD), is important for understanding and conserving biodiversity. Its widely documented negative relationship with species richness has led to the view that species-poor communities may be disproportionately important for regional diversity. Here, we derive neutral expectations for richness–uniqueness relationships and show that negative slopes can arise from neutral assembly rather than necessarily reflecting an ecological signal of uniqueness. Neutral simulations validate the analytical expectations across abundance- and incidence-based LCBD formulations. Applying the framework to the Barro Colorado Island tropical forest and North American Breeding Bird Survey, we show that accounting for neutral sampling expectations substantially changes the ranking and spatial distribution of ecologically unique sites. By separating neutral sampling expectations from excess uniqueness, our framework provides a principled basis for interpreting ecological uniqueness and informing biodiversity conservation.

## 1. Introduction

Ecological uniqueness describes the extent to which a local community differs from other communities within a region and contributes to regional variation in species composition (Lu *et al*. 2007; Legendre & De Cáceres 2013). Local contribution to beta diversity (LCBD) provides a widely used measure of uniqueness that quantifies each site’s share of total compositional variation (Legendre & De Cáceres 2013). Because compositionally distinctive communities may harbor species or assemblages that are poorly represented elsewhere, LCBD has increasingly been used in conservation prioritization to identify sites that richness-based criteria may overlook (Dubois *et al*. 2020; Hill *et al*. 2021; Heino *et al*. 2022; Duflot & Vähätalo 2024).

A striking and pervasive empirical pattern is that ecological uniqueness declines with local species richness. Negative LCBD–richness relationships occur across plants, birds, and insects, for both incidence- and abundance-based data (da Silva *et al*. 2018; Dubois *et al*. 2020; García-Navas *et al*. 2022; Luukkonen *et al*. 2024), and predominate globally (Chen *et al*. 2026). This creates an apparent conservation dilemma: because species-poor communities frequently exhibit the highest LCBD values, prioritizing species richness alone may overlook distinctive assemblages (Dubois *et al*. 2020; Dansereau *et al*. 2022). Accordingly, studies have advocated jointly considering richness and uniqueness in conservation planning (Hill *et al*. 2021; Chen *et al*. 2026). Yet this recommendation rests on an assumption that has rarely been scrutinized: that high LCBD at species-poor sites reflects genuine ecological distinctiveness rather than an expectation generated by stochastic community assembly and sampling.

What would make this assumption valid? Prevailing explanations attribute the negative LCBD–richness relationship to deterministic mechanisms that render species-poor sites compositionally distinctive. Environmental filtering offers one route: rare or extreme habitats support restricted subsets of the regional species pool, yielding assemblages that are simultaneously species-poor and distinctive (Qiao *et al*. 2015; Yao *et al*. 2021; Schwesig *et al*. 2025). Spatial isolation provides another, as weakly connected sites, such as headwater streams, experience less compositional exchange (de Oliveira *et al*. 2020; Li *et al*. 2020). Under these mechanisms, high LCBD at species-poor sites can represent a genuine ecological signal whose conservation may protect compositional information missed by richness-based priorities (Hill *et al*. 2021).

Recent work has shown, however, that LCBD–environment relationships can depend on how environmental gradients are sampled because directional turnover affects the pairwise dissimilarities underlying LCBD (Hernández-Carrasco *et al*. 2026). This raises a more fundamental possibility: negative LCBD–richness relationships may emerge, at least in part, without ecological differentiation among sites. This possibility has a natural foundation in neutral theory, in which the regional species abundance distribution, stochastic dispersal, and demographic drift jointly shape species distributions among local communities (Hubbell 2001). Because the same neutral parameters determine both local richness (Etienne & Alonso 2005; Rosindell & Cornell 2007; Rosindell & Chisholm 2021) and among-sites beta diversity (Chave & Leigh 2002; Condit *et al*. 2002; Xing 2026), richness and uniqueness can be coupled outcomes of a single neutral assembly process rather than independent community properties. Reported effects of species-pool and dispersal attributes on empirical LCBD–richness relationships (Chen *et al*. 2026) may likewise arise partly through this coupling. Whether neutral expectations are quantitatively sufficient to account for observed patterns remains untested, however, because no richness-matched analytical baseline for LCBD has been derived.

Here, we develop such a framework from neutral theory. Starting from a classical neutral birth–death–immigration process (Kendall 1949), we derive exact, closed-form expectations for both abundance- and incidence-based LCBD. Specifically, we formulate three neutral baselines, a parameterized model describing the relationship between expected LCBD and expected richness, and two conditional models predicting expected LCBD at observed local richness under fixed versus stochastic regional species abundances. We validate these expectations with neutral simulations and apply them to the Barro Colorado Island (BCI) tropical forest and the North American Breeding Bird Survey (BBS). Finally, we quantify LCBD deviation (ΔLCBD) as departure from the conditional neutral expectation and show that accounting for neutral sampling substantially changes ecological interpretation and conservation prioritization.

## 2. Materials and Methods

### 2.1 Theory

#### 2.1.1 Neutral theory and local species richness

We developed neutral expectations for local species richness and ecological uniqueness by treating local communities as samples from a shared regional metacommunity. Neutral theory has generated extensive predictions for local richness and community composition (Rosindell *et al*. 2011; Rosindell & Chisholm 2021). Here we briefly summarize the components required for deriving the richness–uniqueness relationship.

At the regional scale, neutral theory predicts a Fisher log-series species-abundance distribution (SAD). The probability that a randomly selected species has regional abundance *n* ≥ 1 is

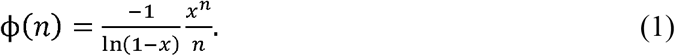

This distribution arises from neutral birth–death–immigration models and is the analytical limit of neutral metacommunity dynamics under speciation and demographic drift (Volkov *et al*. 2003). Parameter *x* characterizes the regional SAD and can be interpreted as an effective regional birth-to-death ratio.

At the local scale, the Kendall birth–death–immigration process predicts a stationary negative-binomial distribution (NBD), whose local absence probability depends on expected abundance *μ*_*n*_ and aggregation parameter *k*_*n*_. These are related through the local birth-to-death ratio 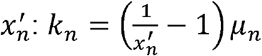 (Kendall 1949). Under strict neutrality, species share local birth and death rates, so *k*_*n*_ scales linearly with expected local abundance. We therefore write *k*_*n*_ =*cμ*_*n*_, where *c* controls local aggregation (He & Legendre 2002).

For an NBD with mean *μ*_*n*_ and aggregation parameter *k*_*n*_, the probability that a species occurs locally is: 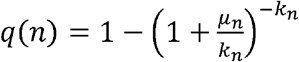 . Because 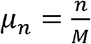, where *M* is the number of local communities, *k*_*n*_ = *cμ*_*n*_ gives:

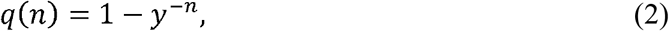

where 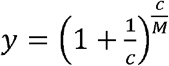 summarize local spatial structure generated by the neutral process. Expected local richness is the sum of occurrence probabilities across the regional species pool of *S* species. Substituting eqns (1) and (2) gives

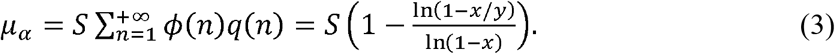

#### 2.1.2 LCBD and site-level uniqueness

We considered an abundance-based coverage-deficit framework and an incidence-based Sørensen framework. Because they weight compositional differences differently, their neutral expectations need not respond identically to richness conditioning.

The abundance framework defines beta diversity as the abundance-weighted local species absence (Xing 2026):

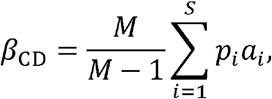

where *p*_*i*_ is regional relative abundance and *a*_*i*_ the probability that species *i* is locally absent. With equal regional abundances, this formulation reduces to classical Whittaker multiplicative beta diversity and therefore generalizes the same conceptual measure to unequal species abundances.

Importantly, this index partitions additively among sites: 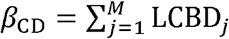(Xing 2026), where

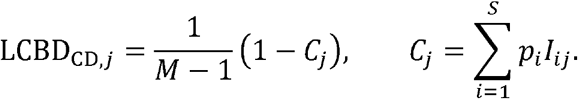

Here *I*_*ij*_ indicates occurrence of species *i* at site *j. C*_*j*_ is the Good-Turing sample coverage of site *j* relative to the regional pool, so 1-C_*j*_ is the regional abundance mass not represented at that site. Coverage deficit therefore provides a direct abundance-weighted measure of site uniqueness.

For incidence data, we defined focal-site uniqueness as its mean Sørensen dissimilarity to the other *M*− 1 sites from the same regional community. For sites *X* and *Y*, let 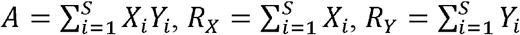, where *X*_*i*_,*Y*_*i*_ ∈{0,1} denote species occurrence. Sørensen dissimilarity is 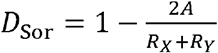. Mean pairwise dissimilarity is an exact positive affine transformation of variance-normalized LCBD and therefore yields identical within-region site rankings (Legendre & De Cáceres 2013; Baselga & Gómez-Rodríguez 2026).

Because partitioned site-level LCBD mechanically decreases with the number of sampled sites, we report uniqueness after multiplication by *M*. Coverage-deficit uniqueness is therefore 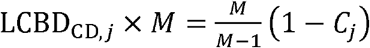, and incidence-based uniqueness is 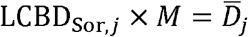, where 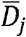 is mean Sørensen dissimilarity between site *j* and the other *M*−1 sites. This rescaling removes the 1/*M* dependence of site-level partitioning while preserving rankings.

#### 2.1.3 Matching mean richness: the parameterized baseline

Our first neutral reference asks how expected uniqueness changes across neutral regimes with different *expected* local richness. For a specified richness *α*, we set *μ*_*α*_ = *α* in eqn (3) and adjust *y* until the model has the desired expectation. The resulting parameterized baseline, *B*^par^(*α*), therefore compares neutral regimes at fixed *x*.

For coverage-deficit, integrating the abundance-weighted absence probability over the Fisher SAD and eliminating *y* using eqn (3) gives

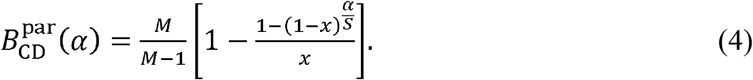

For Sørensen uniqueness, let 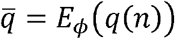 and 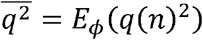. Under richness matching, 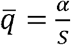, while 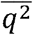 is evaluated at the corresponding *y* implied by eqn (3). Defining 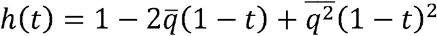, the exact parameterized expectation is

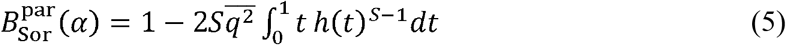

Eqns (4) and (5) demonstrate that expected uniqueness declines systematically with expected local richness under neutral assembly (Fig. 1a-b). These parameterized baselines describe the structural coupling between mean richness and mean uniqueness implied by neutral assembly. It should not, however, be confused with the expected uniqueness of sites that *realize* different richness values within a given neutral system.

**Figure 1.**
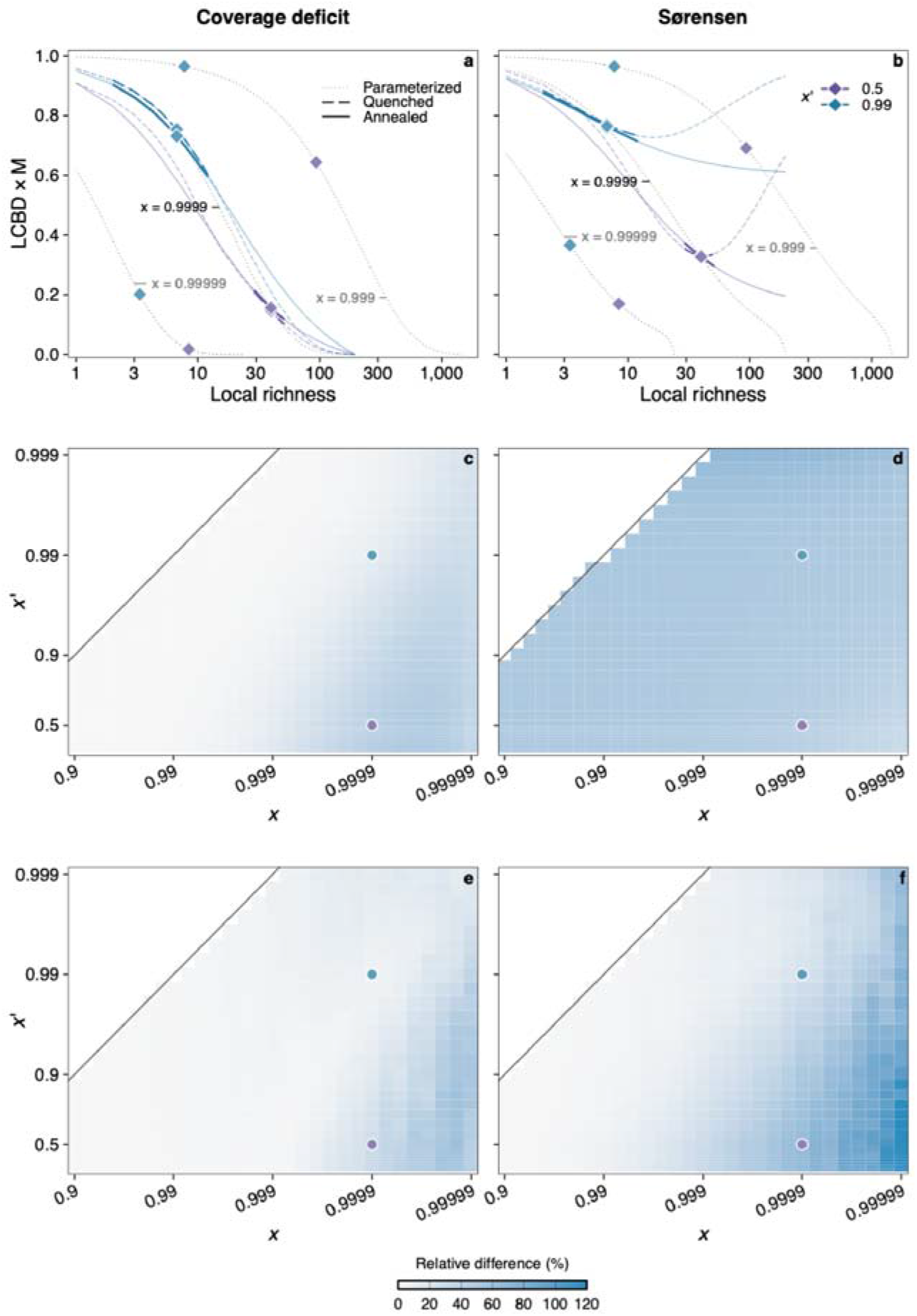
Comparison of three neutral richness–uniqueness baselines for two beta-diversity metrics. (a, b) Expected relationships for *N* = 211,250 individuals and *M* = 1,250 local communities. Parameterized relationships are shown for different *x*, and conditional expectations for *x* = 0.9999 and different local birth-to-death ratios *x’*. Diamonds mark mean expected local richness with thickened segments indicate the 95% likely range of realized local richness. (c, d) Relative slope difference between the annealed and parameterized expectations, **100|*b*_*A*_ −*b*_*p*_|/|*b*_*p*_|**. (e, f) Relative slope difference between the annealed and quenched expectations, **100|*b*_*A*_ −*b* _*Q*_ /|*b*_*A*_ |**, summarized as the median across 100 regional SAD realizations. Circles mark the parameter combinations illustrated in panels a and b.

#### 2.1.4 Conditioning on realized richness: quenched and annealed baselines

Conditioning on observed richness *α* differs fundamentally from matching expected richness. We distinguish two conditional baselines according to whether the regional abundance configuration is treated as fixed or stochastic.

##### Quenched conditional baseline

The quenched baseline (*B*^*Q*^) fixes one realized regional abundance vector *n* = (*n*_1_,…,*n*_*s*_) and the local occurrence structure *y*. Let 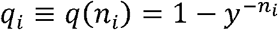 and 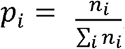. Local richness is *R*=∑_*i*_ *X*_*i*_, where *X*_*i*_ is a Bernoulli occurrence indicator with probability qi, and let *R* _−*i*_ =∑_*k* ≠*i*_ *X*_*k*_ be the richness contributed by all species other than species *i*. Conditional on *R* = *α*, the probability that species *i* is present is

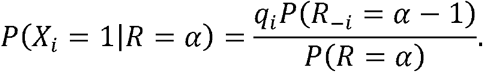

The exact coverage-deficit expectation is therefore:

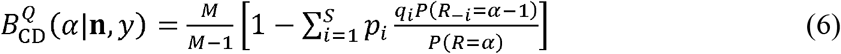

Eqn (6) gives expected regional abundance coverage of a site realizing richness *α*, conditional on the particular realized regional abundance configuration. Richness probabilities follow a Poisson-binomial distribution determined by the species-specific *q*_*i*_ (Appendix S1).

For Sørensen uniqueness, the focal site is conditioned on *R*_*X*_= α, whereas the comparison site samples independently from the same fixed regional pool. The exact quenched baseline is:

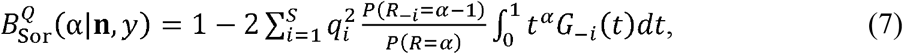

where *G* _−*i*_ = Π_*k* ≠*i*_(1−*q*_*k*_ +*q*_*k*_*t*) is the probability-generating function for comparison-site richness excluding species *i*. The integral averages the Sørensen denominator over the full richness distribution of the comparison site. Equivalent probability and finite-sum forms, together with numerically stable recursions, are provided in Appendix S1.

##### Annealed conditional baseline

The annealed baseline (*B*^*A*^) fixes the macroecological parameters (*x,y*) rather than a realized regional SAD. Regional abundances therefore remain stochastic under the Fisher log-series and are averaged over after conditioning on *R*_*X*_= α

Conditioning on species presence or absence in the focal site yields posterior regional-abundance distributions with probability-generating functions 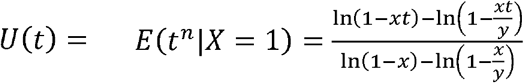, and 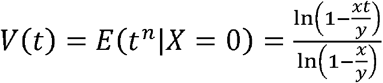. For coverage deficit, the annealed expectation is:

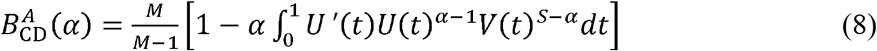

The same posterior distributions determine Sørensen uniqueness. Because local absence for a species of regional abundance *n* is *y*^−*n*^, we have 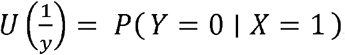, and 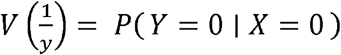, for an independently sampled comparison site *Y*. The annealed Sørensen expectation is therefore:

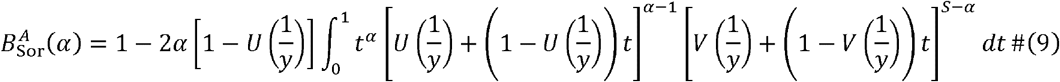

Thus, both annealed baselines are determined by posterior abundance distributions *U* and *V*. For fixed *S* and *M*, their unknown macroecological parameters are *x* and *y*.

The quenched and annealed baselines differ only in how regional stochasticity is treated. The quenched baseline conditions on one realized regional abundance configuration and averages over local assembly, whereas the annealed baseline additionally averages over regional SAD realizations consistent with the same macroecological parameters. Both are exact conditional expectations but answer different inferential questions.

#### 2.1.5 Interpretation, choice of baseline, and excess uniqueness

The three baselines answer related but distinct questions. The parameterized baseline describes how mean richness and mean uniqueness covary across differently parameterized neutral systems. Conditional baselines instead ask how unique a site is expected given realized richness *α* under specified neutral parameters. Consequently, *B*^par^ (*α*) is not generally equal to either *B*^*Q*^ (*α*) or *B*^*A*^(*α*). Around the high-probability range of local richness, conditional relationships are typically shallower than the corresponding parameterized relationship, especially for Sørensen uniqueness (Fig. 1c-d). We therefore use the parameterized baseline to characterize structural richness– uniqueness coupling and conditional expectations to evaluate empirical deviations.

Quenched and annealed conditional baselines differ in whether the realized regional abundance configuration is fixed. Their practical difference depends on local-regional separation. When local communities are relatively autonomous from the regional pool, the two expectations converge over the likely richness range; when local composition depends strongly on continual regional immigration, they can diverge substantially (Fig. 1e, f). Because regional stochasticity is intrinsic to neutral dynamics and empirical regional pools are rarely literally fixed, we recommend the annealed baseline as the default for empirical applications; the quenched baseline is appropriate when inference explicitly conditions on a realized regional configuration.

The distinction is particularly important for Sørensen uniqueness. Under quenched conditioning, increasing focal richness does not simultaneously increase expected comparison-site richness. Shared richness therefore becomes constrained by the comparison site while the Sørensen denominator continues to increase, allowing the quenched expectation to reach a minimum and then rise at high focal richness. This upturn is absent from the annealed expectation because conditioning on focal richness also updates the expected regional state and hence comparison-site richness and composition. Coverage deficit is less sensitive because it is a linear function of the regional abundance mass represented locally, so quenched and annealed expectations are generally more similar (Fig. 1).

We define excess uniqueness for site *j* as

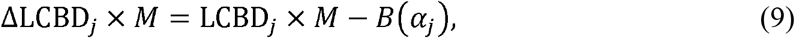

where *α*_*j*_ is observed richness and *B*(*α*_*j*_) is the fitted conditional expectation. Positive ΔLCBD indicates greater uniqueness than expected at observed richness; negative values indicate lower-than-expected uniqueness.

### 2.2 Validation and empirical applications

#### 2.2.1 Neutral simulations

We evaluated the analytical models using spatially explicit neutral simulations retaining distance-dependent dispersal and spatially implicit simulations in which local communities are conditionally independent given regional abundances. Spatially explicit communities were generated with rcoalescence (Thompson *et al*. 2020). We simulated a closed 650×325-cell landscape, with one individual per cell, rescaled to 1,000 m × 500 m. The speciation rate (1.77 × 10□ □) and Gaussian dispersal-kernel standard deviation (σ= 26 cells) represented a dispersal-limited neutral forest broadly consistent with previous BCI applications (Chisholm 2024), while yielding stem density and regional richness close to those of the 50-ha plot. The resulting community contained 211,250 individuals and 306 species. We aggregated the landscape at 5 m × 5 m, 20 m × 20 m and 100 m × 100 m, corresponding to *M* = 20,000, 1,250 and 50 local communities, respectively.

To generate spatially implicit counterparts with the same realized regional abundance structure, we retained the regional SAD from the spatially explicit simulation. At each grain, we estimated local structure parameter *c* (equivalently 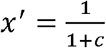) from the spatially explicit communities by composite marginal likelihood under the NBD model with *k*_*n*_ =*cμ*_*n*_ (Xing 2026). Estimated *x’*^′^ values were 0.009, 0.206 and 0.863. For each species *i*, local abundances across *M* communities were then drawn independently from an NBD with mean *N*_*i*_/*M* and size *k*_*i*_ = *cN*_*i*_/*M*, where *N*_*i*_ is its realized regional abundance.

We separately validated annealed conditional baselines (eqns 8 and 9) against Monte Carlo conditional expectations. For each of *x’* = 0.5 and 0.99, we generated 10,000 community matrices. For each matrix, a regional SAD was first drawn from a Fisher log-series distribution with *S* = 300 and *x* = 0.9999, followed by local NBD sampling across *M* =1,250 communities. Uniqueness values were pooled by realized integer richness and averaged to obtain the Monte Carlo conditional expectation. Agreement with analytical expectations was quantified by frequency-weighted signed bias and root mean squared error (RMSE).

#### 2.2.2 Empirical data

The BCI 50-ha forest dynamics plot is located in lowland tropical moist forest in central Panama. All free-standing woody stems with diameter at breast height ≥ 1 cm are tagged, mapped, measured and identified following standardized protocols (Condit *et al*. 2019). We analyzed 221,527 living individuals of 299 species in the 2015 census. Main analyses used 20 m × 20 m quadrats; 5 m × 5 m and 100 m × 100 m grains were additionally examined to evaluate grain dependence. Quadrats were assigned to the habitat types of Harms *et al*. (2001).

The BBS surveys birds annually along standardized 39.2-km roadside routes comprising 50 stops (Sauer *et al*. 2017; Ziolkowski *et al*. 2025). We used the 2024 surveys from the 2025 release, retaining standard routes (RouteTypeID = 1,

RouteTypeDetailID = 1, RPID = 101). Counts were summed across stops to obtain route-level species abundances. Routes were treated as local communities and Bird Conservation Regions (BCRs) as regional communities. We retained 2,484 routes from 28 BCRs with at least 20 sampled routes and used the Northern Rockies (BCR 10; 161 routes) as a worked example.

Within each BCR, conservation representation was evaluated using Partners in Flight (PIF) regionally important species (RI = 1 in the Avian Conservation Assessment Database; Panjabi *et al*. 2021) and U.S. Fish and Wildlife Service breeding Birds of Conservation Concern (U.S. Fish and Wildlife Service 2021). Existing protection was assessed from the March 2026 World Database on Protected Areas and Conserved Areas (UNEP-WCMC & IUCN 2026). We retained designated or established polygons, dissolved overlaps and classified a route as protected if its starting point fell within a protected polygon.

#### 2.2.3 Model fitting and statistical analysis

For spatially implicit and explicit simulations, we calculated coverage-deficit and Sørensen uniqueness at all three grains. Parameterized (eqns 4 and 5) and quenched conditional (eqns 6 and 7) baselines were fitted separately to each simulated community set by bounded optimization minimizing mean squared difference between observed site-level uniqueness and analytical expectation. ΔLCBD was calculated as observed LCBD minus the corresponding quenched expectation. We then tested whether ΔLCBD retained richness dependence under neutral assembly. Spatially implicit expectations were additionally evaluated using the known generating parameters.

For empirical analyses, coverage-deficit and Sørensen uniqueness were analyzed separately. Parameterized and annealed baselines were fitted independently for each BCI grain and BBS BCR using the same optimization procedure. ΔLCBD was calculated by subtracting the fitted annealed expectation at observed richness from observed LCBD.

To examine the consequences for conservation prioritization, we compared local communities in the top 5% of the raw-LCBD and by ΔLCBD rankings. For BCI, we compared selected quadrats spatially and by representation of the regional pool and rare species. Rare species were defined as those represented by fewer than one individual per hectare, following Hubbell (2013). For BBS, comparisons within each BCR considered representation of the regional species pool, PIF regionally important species, USFWS Birds of Conservation Concern, and protection status. Alternative selection thresholds were examined for sensitivity.

## 3. Results

### 3.1 Numerical validation

In spatially implicit simulations, parameterized baselines fitted separately at the three grains were virtually indistinguishable (Fig. 2a, c). Quenched conditional expectations closely reproduced the richness–uniqueness relationships under both frameworks, and ΔLCBD showed no systematic relationship with richness at any grain (Fig. 2e, g).

**Figure 2.**
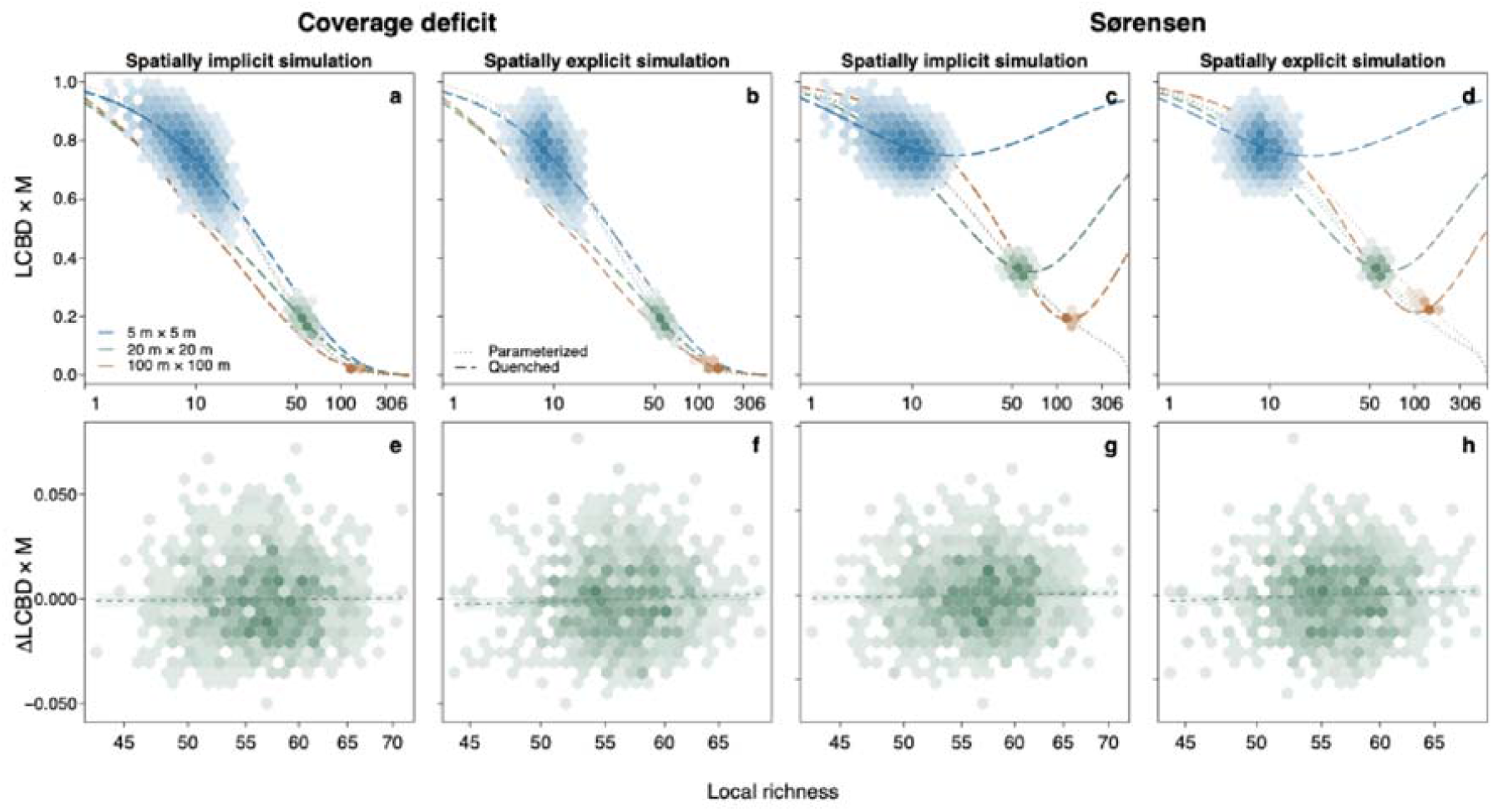
Validation of the quenched conditional baselines under spatially implicit and spatially explicit neutral assembly. (a–d) Relationships between uniqueness and local richness across three spatial grains for one realized regional SAD. Hexagon shading indicates site density. (e–h) Relationships between ΔLCBD from the quenched expectations and local richness at the 20 m × 20 m grain, with dashed lines indicate nonsignificant regressions (*P* > 0.05).

The conditional expectations remained effective under spatially explicit neutral assembly, although dispersal limitation introduced grain dependence absent from implicit simulations. Parameterized baselines fitted at one grain did not transfer across grains, whereas grain-specific quenched expectations broadly tracked observed relationships except for Sørensen uniqueness at 100 m × 100 m grain (Fig. 2b, d). This departure indicates that distance-dependent spatial structure can generate residual richness dependence not captured by the spatially implicit baseline. Post-hoc spatial autoregressive analyses substantially reduced this residual relationship (not shown).

Annealed analytical expectations closely matched Monte Carlo conditional means (Fig. 3). Across supported richness ranges and contrasting levels of local aggregation, both RMSE and absolute signed bias were below 5×10^-4^ for coverage-deficit and Sørensen uniqueness.

**Figure 3.**
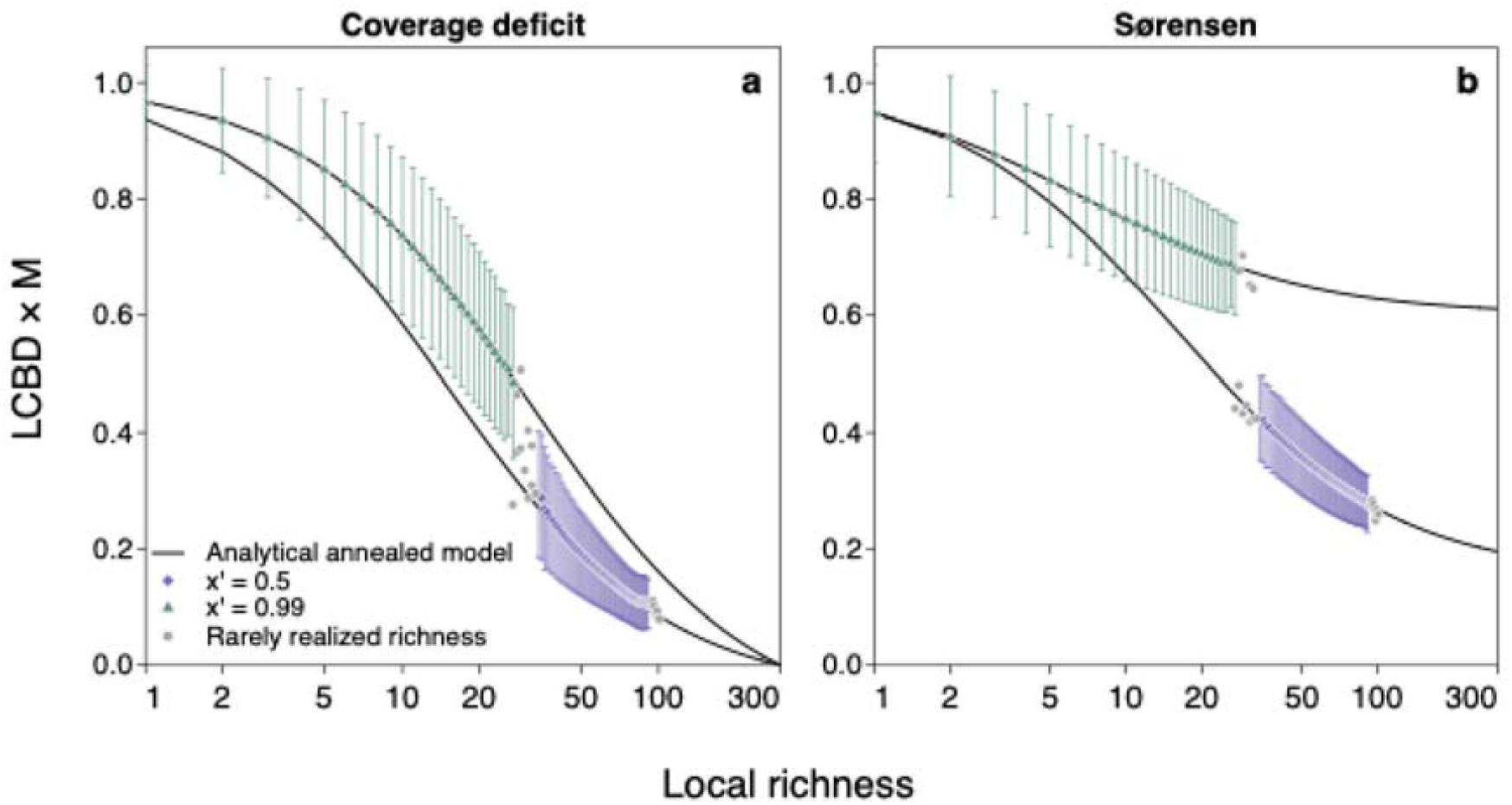
Monte Carlo validation of the annealed conditional baseline. Analytical expectations are compared with Monte Carlo conditional means for *S* = 300, *M* = 1,250, *x* = 0.9999, and *x’* = 0.5 or 0.99. Points and error bars show mean ±1.96 SD of uniqueness at each realized integer richness, pooled across 10,000 independently generated community matrices. Grey points denote richness values realized in fewer than 50 matrices.

### 3.2 BCI tropical forest

In BCI, coverage-deficit uniqueness declined with richness at all three grains. Annealed baselines closely tracked these relationships, whereas parameterized baselines fitted separately at the three grains diverged substantially (Fig. 4a). At 20 m × 20 m, the top 5% of quadrats ranked by ΔLCBD represented 79.3% of the 299 plot species and 46.4% of the 110 rare species, compared with 72.6% and 31.8%, respectively, under raw LCBD (Fig. 4b). The two rankings shared 35 of their 63 selected quadrats but differed spatially. Both included the swamp; ΔLCBD additionally selected clustered low-plateau and streamside quadrats, whereas quadrats selected only by raw LCBD were more widely scattered (Fig. 4c). Results were qualitatively consistent for Sørensen uniqueness (Fig. S1).

**Figure 4.**
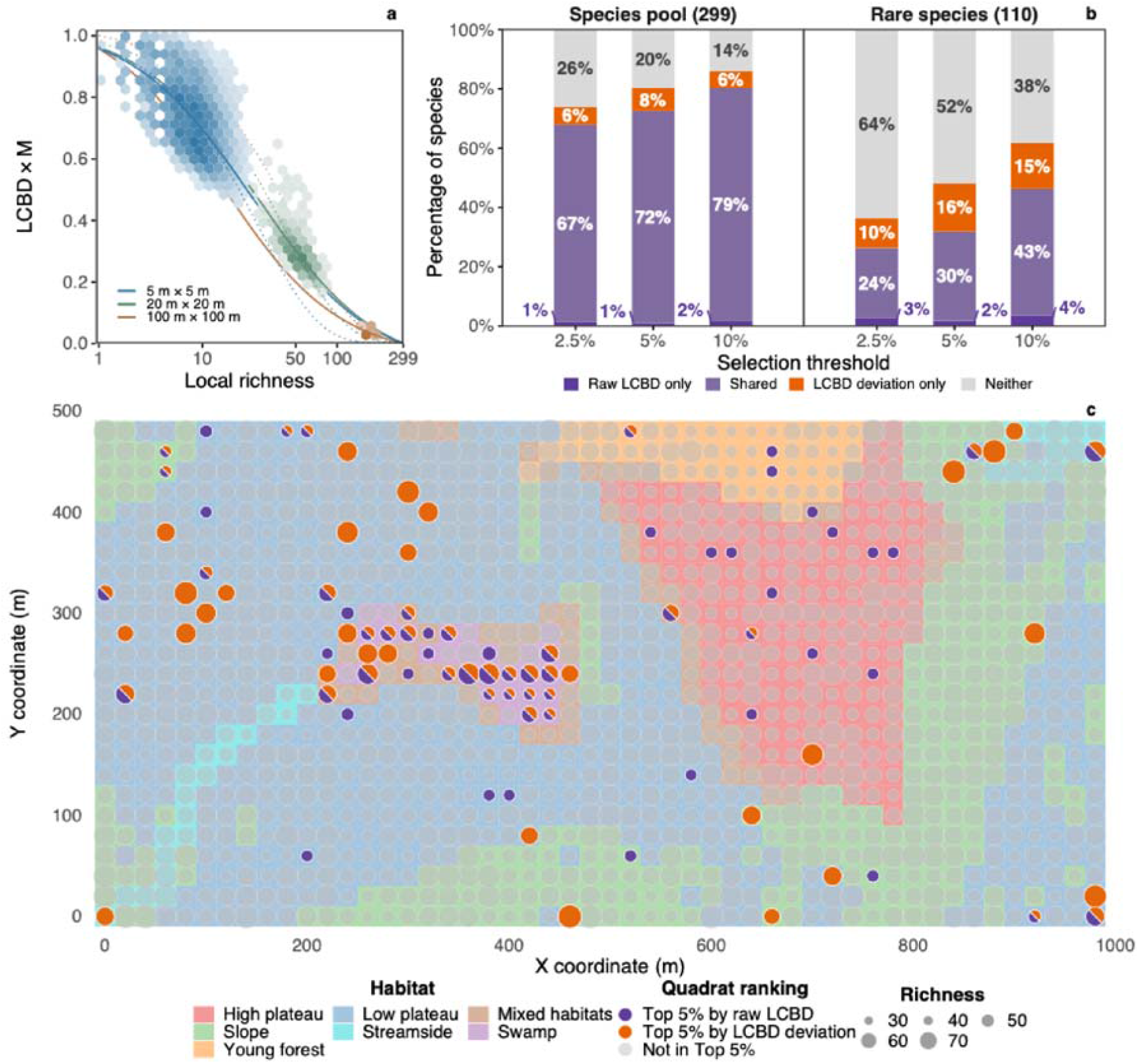
Richness–uniqueness relationships in the BCI forest plot. (a) Coverage-deficit LCBD versus local richness across three spatial grains. Hexagon shading indicates quadrat density; dotted and solid curves show fitted parameterized and annealed conditional expectations, respectively. (b) Representation of the regional species pool and rare species by 20 m ×20 m quadrats selected under raw-LCBD or ΔLCBD rankings at alternative thresholds. (c) Spatial distribution of quadrats in the top 5% of the two rankings at the 20 m × 20 m grain.

### 3.3 North American BBS

Across the 28 BCRs, BCR-specific annealed expectations closely tracked the negative LCBD–richness relationship (Figs. S2, S3), illustrated by the Northern Rockies (Fig. 5a). Top-5% raw-LCBD and ΔLCBD selections comprised 138 routes each, with 79 (57.2%) shared. ΔLCBD selections represented a larger fraction of the regional pool in 22 of 28 BCRs, with median representation of 56.3% versus 45.1% under raw LCBD (Fig. 5b). They also represented more conservation-listed species (Fig. 5c-d; Table S1): the median within-BCR difference was 5.6 percentage points for PIF regionally important species (higher under ΔLCBD in 20 of 28 BCRs) and 9.3 percentage points for USFWS Birds of Conservation Concern (higher in 18 of 26 BCRs with BCC lists). The two rankings showed no consistent difference in the proportion of protected routes. The same qualitative patterns were obtained under the Sørensen framework (Figs. S3, S4; Table S2).

**Figure 5.**
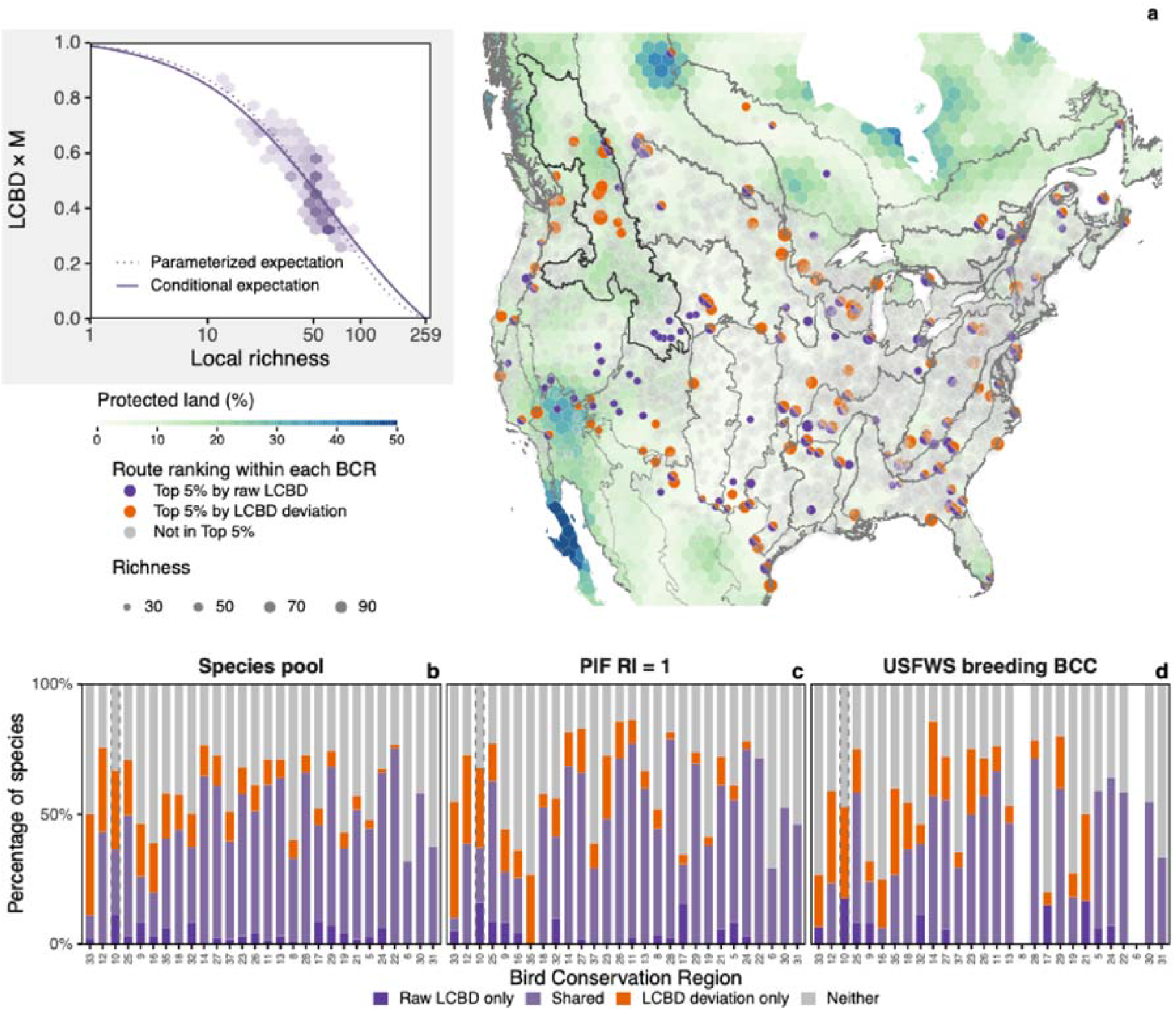
Richness–uniqueness relationships across North American BBS Bird Conservation Regions. (a) BBS routes in the top 5% of raw-LCBD and ΔLCBD rankings within each BCR, shown over protected-land coverage on a 100-km-spaced hexagonal grid. The inset shows coverage-deficit uniqueness versus richness in the Northern Rockies (BCR 10, highlighted across panels). (b–d) Representation of the regional species pool (b), PIF regionally important species (c), and USFWS breeding Birds of Conservation Concern (d) under the two rankings.

## 4. Discussion

### 4.1 A formal baseline for richness-dependent uniqueness

Ecological uniqueness is commonly interpreted as evidence that a local community has been shaped by unusual environmental conditions, dispersal histories or species interactions (da Silva *et al*. 2018; Li *et al*. 2020; Hill *et al*. 2021; Yao *et al*. 2021; Chen *et al*. 2024), yet the tendency for species-poor communities to have high LCBD has long been recognized. In their original formulation of LCBD, Legendre and De Cáceres (2013) reported a negative LCBD–richness correlation, while explicitly noting that it was “not … a general or obligatory relationship”. Our contribution is therefore not to establish this association but to provide a formal expectation for its emergence under neutral assembly.

Richness and uniqueness are coupled outcomes of the same regional abundance structure and local sampling process: richer sites represent more of the regional abundance mass and, for incidence data, tend to share more species with other sites. A negative richness– uniqueness relationship therefore need not imply environmental filtering, spatial isolation, or other deterministic differentiation. In both BCI and BBS, neutral baselines captured much of the observed decline, suggesting that a substantial component of this widespread pattern can arise from neutral sampling alone.

A recent global meta-analysis identified species-pool attributes as the dominant predictors of incidence-based richness–uniqueness relationships, while dispersal limitation was especially important for abundance-based relationships (Chen *et al*., 2026). Our theory provides a mechanistic interpretation while showing that these effects are scale dependent. Holding regional extent, and hence metacommunity size, constant, increasing gamma diversity corresponds to decreasing *x* in the Fisher log-series. When local–regional separation (*x’*) is weak, the model predicts a steeper negative richness– uniqueness relationship with increasing gamma diversity, qualitatively matching the meta-analytic pattern (Fig. 1a, b). At stronger local–regional separation, however, this relationship can weaken or even reverse. Species-pool attributes therefore do not uniquely determine the slope; their effect depends on where local communities sample the regional abundance structure.

This dependence is captured by *x’*, which determines both the typical local-richness range and effective separation between local and regional dynamics. At the weak-separation end, increasing *x’*—consistent with stronger dispersal limitation or greater local demographic autonomy—generally steepens the negative relationship, particularly for coverage-deficit uniqueness (Fig. 1a, b). This prediction is qualitatively consistent with the stronger role of dispersal limitation in abundance-based richness–uniqueness relationships reported by Chen *et al*. (2026). More importantly, our theory clarifies that species-pool structure, spatial grain and dispersal limitation need not act independently: under neutrality their effects are interactive in shaping richness–uniqueness relationships. The direction and strength of these effects are therefore context dependent, and a formal cross-dataset test of the theoretical predictions remains to be done.

### 4.2 Ecological interpretation and conservation implications

Once the neutral sampling expectation is accounted for, ΔLCBD identifies communities that are more or less distinctive than expected at their observed richness, separating excess uniqueness from high LCBD generated by the neutral richness– uniqueness coupling. In BCI, both raw LCBD and ΔLCBD retained the species-poor swamp, a floristically distinctive habitat recognized previously (Harms *et al*. 2001; Legendre & Condit 2019), showing that accounting for the neutral expectation does not remove genuinely distinctive species-poor communities. Beyond this shared signal, raw LCBD highlighted spatially scattered, low-richness quadrats, whereas ΔLCBD revealed spatially coherent clusters of extreme excess uniqueness (Fig. 4c), including streamside quadrats and two low-plateau clusters. Streamside habitats support characteristic species associations (Harms et al. 2001), consistent with the broader role of water availability and drought sensitivity in structuring BCI tree distributions (Comita & Engelbrecht 2009). The low-plateau clusters coincide with distinct subcommunities differing in life-history traits including shade tolerance (Flügge *et al*. 2014). Thus, ΔLCBD can reveal habitat-associated departures from the neutral baseline rather than merely reorder sites. Such patterns are consistent with habitat and disturbance effects, although ΔLCBD itself does not identify mechanism.

Beyond these habitat-associated departures, the pronounced grain dependence of the parameterized richness–uniqueness relationship in BCI (Fig. 4a) suggests that additional departures from the spatially implicit neutral model may also operate across scales. Spatially structured neutral dispersal could contribute to this divergence, but the greater magnitude than in our spatially explicit neutral simulations is also compatible with species-specific departures from the neutral abundance–aggregation relationship. Such departures are plausible in BCI, where rare species experience stronger conspecific negative density dependence than common species(Comita *et al*. 2010), and adult trees show widespread within-species spatial repulsion beyond expectations from stochastic birth, death and limited dispersal (Kalyuzhny *et al*. 2023). These non-neutral demographic processes may therefore modify how aggregation, occupancy and hence richness–uniqueness coupling scale with grain.

Whether high uniqueness reflects genuinely distinctive composition or a sampling-induced expectation also matters for conservation prioritization. Complementarity, a core principle of systematic conservation planning, values sites for the features they add to a regional network (Margules & Pressey 2000; Kukkala & Moilanen 2013). It has motivated the use of LCBD to identify sites of conservation value (Dubois *et al*. 2020; Hill *et al*. 2021). Yet high LCBD can partly arise from the sampling structure, including the neutral richness–uniqueness coupling quantified here and other sampling-dependent properties of compositional dissimilarity (Baselga & Gómez-Rodríguez 2026; Hernández-Carrasco *et al*. 2026). Treating low-richness, high-LCBD sites as intrinsically high-value conservation targets can therefore misdirect site selection. ΔLCBD reduces this risk by quantifying uniqueness relative to its neutral expectation at observed richness. In BCI and most BBS BCRs, ΔLCBD selections represented more of the regional species pool and rare or conservation-listed species than raw-LCBD selections, showing that removing the neutral sampling component can improve regional biodiversity representation by sites identified as compositionally distinctive.

### 4.3 Regional stochasticity in null expectations

Ecological and conservation inferences depend on the baseline against which uniqueness is measured, particularly whether the realized regional state is treated as fixed or stochastic. Many community-ecology null models condition on an observed regional pool, species frequencies, or other properties of the realized community matrix and randomize within that fixed regional context (Ulrich & Gotelli 2010; Chase *et al*. 2011; Kraft *et al*. 2011). Such procedures are analogous to the quenched baseline: inference is conditional on the regional state that happened to be observed. This is appropriate when that state is explicitly the object of conditioning but differs from asking whether a local community is unusual relative to a process capable of generating alternative regional states.

Neutral theory makes the distinction especially clear because stochasticity operates at both regional and local scales. The annealed baseline averages over regional SAD realizations after conditioning on local richness, whereas the quenched baseline fixes one realized regional configuration. Their difference is small when local communities are relatively autonomous from the regional pool but can become substantial under strong continual immigration (Fig. 1). The key distinction is therefore not whether a null model is analytical or simulation based, but which sources of stochasticity defines its null probability space. This conditioning structure is often implicit in community randomizations, yet it directly determines what constitutes a meaningful departure from the null expectation (Xing & He 2021).

### 4.4 Caveats and extensions

Our spatially implicit baselines incorporate abundance-related aggregation through the NBD but not distances among local communities or dispersal kernels generating spatial covariance. The spatially explicit simulations show that this approximation is generally effective at finer grains and for coverage-deficit uniqueness, but can leave residual structure, particularly for Sørensen uniqueness at coarse grain. Future extensions could incorporate spatially explicit neutral theory, although obtaining tractable predictions across spatial scales remains challenging (Etienne & Rosindell 2011; Thompson *et al*. 2020). Empirical analyses can meanwhile use spatial predictors or autoregressive approaches to test and account for residual spatial dependence (Dray *et al*. 2006).

Departures from the neutral baseline need not imply a single deterministic mechanism. They can arise from environmental filtering, species interactions, heterogeneous dispersal, variation in local carrying capacity, or other violations of the assumptions underlying the neutral model. ΔLCBD should therefore be interpreted as excess compositional uniqueness relative to a specified null expectation, not as a direct measure of niche-based assembly. Its ecological value lies in providing a response variable that can subsequently be related to environmental, spatial or trait predictors to evaluate candidate mechanisms. More generally, the analytical strategy developed here is not restricted to strict neutrality. The present model uses the neutral abundance– aggregation relationship *k* = *cμ*, but the same framework could be extended by relaxing this scaling or by replacing the neutral regional and local abundance models with alternatives representing non-neutral assembly. Comparing observed uniqueness against a hierarchy of increasingly structured baselines could help quantify the relative importance of stochastic and deterministic assembly.

## Acknowledgements

We thank the many field workers and data managers who contributed to the collection and curation of the long-term forest dynamics data on Barro Colorado Island (BCI). We also thank the thousands of U.S. and Canadian participants and coordinators whose efforts made the BBS data available. We used ChatGPT (OpenAI) to assist with language editing and code development. All AI-assisted outputs were critically reviewed, revised, and verified by the authors, who take full responsibility for the final content. This work was supported by the National Natural Science Foundation of China (32471623 and 32101280), the Fundamental and Interdisciplinary Disciplines Breakthrough Plan of the Ministry of Education of China (JYB2025XDXM904), and the Innovation Program of the Shanghai Municipal Education Commission (2023ZKZD36).

