## Appendix S1 for "A sampling-theory framework for richness–uniqueness relationships"

### **Appendix S1. Mathematical derivation of the neutral richness–uniqueness baselines**

This appendix derives the three neutral richness–uniqueness baselines introduced in Section 2.1: the parameterized baseline, the quenched conditional baseline, and the Fisher-averaged annealed conditional baseline. Each is derived for both coverage-deficit and Sørensen uniqueness.

The three baselines differ in what is held fixed. The parameterized baseline compares neutral systems with different *expected* *local* *richness*. The quenched baseline conditions on *realized* *richness* while holding one realized regional species-abundance distribution fixed. The annealed baseline also conditions on *realized* *richness* but averages over regional abundance realizations drawn from the same stationary Fisher distribution. These distinctions are central to the derivations below.

Throughout, uniqueness is reported on the $\text{LCBD}\times M$ scale used in the main text, with LCBD’s sum to total beta diversity as used in many empirical applications (for cases where LCBD’s sum to 1 rather than to beta diversity, the two LCBD definitions differ merely by an affine transformation, see section S1.3.2 below). While equations are numbered indecently in this appendix, the equations corresponding to those presented in the main text are put within boxes.

#### **S1.1 Neutral regional and local community model**

Consider a regional species pool containing $S$ species and $M$ equivalent local sampling units. To distinguish random regional abundance from a particular realization, let $N_{i}$ denote the regional abundance of species $i$ as a random variable and $n_{i}$ its realized value.

Under the Fisher ensemble, regional species abundances are independent draws from the log-series distribution

$$\boxed{\phi\left( n \right)=-\frac{x^{n}}{n\ln\left( 1-x \right)}, n=1,2,\ldots,} \left( S1 \right)$$

where $0<x<1$. The probability-generating function of regional abundance is therefore

$$\Psi\left( t \right)=E_{\phi}\left( t^{N} \right)=\frac{\ln\left( 1-xt \right)}{\ln\left( 1-x \right)}. \left( S2 \right)$$

Its mean is

$$E_{\phi}\left( N \right)=-\frac{x}{\left( 1-x \right)\ln\left( 1-x \right)}. \left( S3 \right)$$

At the local scale, a species with regional abundance $n$ has expected local abundance

$$\mu_{n}=\frac{n}{M}.$$

Under the strict-neutral aggregation relationship $k_{n}=c\mu_{n}$, its local abundance follows a negative-binomial distribution with absence probability

$$P\left( X=0\mid n \right)=\left( 1+\frac{1}{c} \right)^{-cn/M}=y^{-n},$$

where

$$y=\left( 1+\frac{1}{c} \right)^{c/M}. \left( S4 \right)$$

The corresponding occurrence probability is

$$\boxed{q\left( n \right)=P\left( X=1\mid n \right)=1-y^{-n}.} \left( S5 \right)$$

For a fixed realized regional pool $\mathbf{n}=\left( n_{1},\ldots,n_{S} \right)$, we write

$$q_{i}\equiv q\left( n_{i} \right)=1-y^{-n_{i}}.$$

Conditional on the regional abundances, species occurrences are assumed independent across species and across local sites. Thus, for two local sites $X$ and $Y$,

$$X_{i},Y_{i}\mid n_{i}\overset{\mathrm{ind}}{\sim}\mathrm{Bernoulli}\left( q_{i} \right). \left( S6 \right)$$

This is an independent-sampling model conditional on regional abundance. The local draws have expected abundance $n_{i}/M$, but abundances across the $M$ sites are not constrained to sum exactly to $n_{i}$.

Because $c>0$,

$$0<c\ln\left( 1+1/c \right)<1,$$

and hence the strict-neutral construction implies

$$1<y<e^{1/M}. \left( S7 \right)$$

This restriction is relevant when interpreting the parameterized richness-matched curves below.

#### **S1.2 Expected richness and the richness bridge**

Averaging the occurrence probability in eqn (S5) over the Fisher distribution gives the marginal occurrence probability of a randomly selected species,

$$\bar{q}=E_{\phi}\left[ q\left( N \right) \right]=1-\Psi\left( 1/y \right)=1-\frac{\ln\left( 1-x/y \right)}{\ln\left( 1-x \right)}. \left( S8 \right)$$

Expected local richness is therefore

$$\boxed{E\left( R \right)=S\bar{q}=S\left[ 1-\frac{\ln\left( 1-x/y \right)}{\ln\left( 1-x \right)} \right],} \left( S9 \right)$$

which is eqn (3) of the main text.

For the parameterized baseline, expected richness is matched to a specified value $\alpha$. Setting $E\left( R \right)=\alpha$ gives

$$\frac{\ln\left( 1-x/y \right)}{\ln\left( 1-x \right)}=1-\frac{\alpha}{S},$$

and hence

$$1-\frac{x}{y}=\left( 1-x \right)^{1-\alpha/S},$$

or

$$\frac{x}{y}=1-\left( 1-x \right)^{1-\alpha/S} \left( S10 \right)$$

Equation (S10) is the richness bridge used to eliminate $y$ from the parameterized baselines.

A technical distinction is useful here. Equation (S10) formally assigns a $y$ to any $0<\alpha<S$, whereas the strict-neutral relationship in eqn (S4) restricts $y$ to the interval in eqn (S7). The physically attainable expected-richness range at fixed $x$, $S$, and $M$ is therefore narrower than $\left( 0,S \right)$. Richness-matched curves outside that range should be interpreted as a formal continuation of the analytical relationship rather than as values generated by varying $c>0$ within the strict-neutral model. This distinction does not affect the conditional models, for which $x$ and $y$ remain fixed and every integer richness from 0 to $S$ has positive probability whenever $0<q_{i}<1$.

#### **S1.3 Site-level uniqueness measures**

##### **S1.3.1 Coverage deficit**

For a realized regional pool, define regional relative abundance

$$p_{i}=\frac{n_{i}}{\sum_{k=1}^{S} n_{k}}.$$

For a focal site $X$, regional abundance coverage is

$$C_{X}=\sum_{i=1}^{S} p_{i}X_{i}. \left( S11 \right)$$

The corresponding scaled coverage-deficit uniqueness is

$$U_{\mathrm{CD}}\left( X \right)=\frac{M}{M-1}\left( 1-C_{X} \right). \left( S12 \right)$$

When all species have equal regional abundance and mean site richness is $\bar{\alpha}$,

$$U_{\mathrm{CD}}=\frac{M}{M-1}\left( 1-\frac{\bar{\alpha}}{S} \right). \left( S13 \right)$$

Writing Whittaker multiplicative beta diversity as

$$\beta_{W}=\frac{S}{\bar{\alpha}},$$

eqn (S13) becomes

$$U_{\mathrm{CD}}=\frac{M}{M-1}\left( 1-\frac{1}{\beta_{W}} \right). \left( S14 \right)$$

Thus, under equal species abundances, coverage deficit is a monotonic transformation of Whittaker multiplicative beta.

The quantity $C_{X}$ is the population-level abundance coverage of the focal site relative to the regional pool. It is conceptually related to Good–Turing sample coverage, but because regional relative abundances $p_{i}$ are treated here as known model quantities, it should not be confused with a coverage estimator constructed from singleton frequencies.

##### **S1.3.2 Sørensen uniqueness and its relation to LCBD**

For two nonempty sites $X$ and $Y$, define

$$R_{X}=\sum_{i} X_{i}, R_{Y}=\sum_{i} Y_{i}, A=\sum_{i} X_{i}Y_{i}.$$

Sørensen dissimilarity is

$$D_{\mathrm{Sor}}\left( X,Y \right)=1-\frac{2A}{R_{X}+R_{Y}}. \left( S15 \right)$$

For focal site $j$, the operational incidence-based uniqueness score used in the main text is

$$U_{Sor,j}=\bar{D}_{j}, \left( S16 \right)$$

where

$$\bar{D}_{j}=\frac{1}{M-1}\sum_{\mathcal{l\neq}j} D_{j\mathcal{l}}.$$

This score preserves the rankings of variance-normalized LCBD when LCBD is calculated from square-root Sørensen distances. To show this, let the distance supplied to the variance decomposition be

$$d_{j\mathcal{l}}=\sqrt{D_{j\mathcal{l}}},$$

so that $d_{j\mathcal{l}}^{2}=D_{j\mathcal{l}}$. Define

$$r_{j}=\sum_{\mathcal{l=}1}^{M} D_{j\mathcal{l}}, T=\sum_{j=1}^{M} \sum_{\mathcal{l=}1}^{M} D_{j\mathcal{l}}.$$

Double centering of the squared-distance matrix gives the site sum of squares

$$g_{jj}=\frac{r_{j}}{M}-\frac{T}{2M^{2}},$$

while total beta variance is

$$\sum_{j} g_{jj}=\frac{T}{2M}.$$

Variance-normalized LCBD is therefore

$$LCBD_{j}^{\mathrm{var}}=\frac{2r_{j}}{T}-\frac{1}{M}=\frac{2\left( M-1 \right)}{T}\bar{D}_{j}-\frac{1}{M}. \left( S17 \right)$$

Within a dataset, eqn (S17) is a positive affine transformation of $\bar{D}_{j}$, and the two measures therefore produce identical site rankings.

This exact relation does not hold if raw Sørensen dissimilarity itself is supplied as the distance to the variance decomposition, because the squared distances would then be $D_{\mathrm{Sor}}^{2}$.

All conditional Sørensen results below are given for $R_{X}=\alpha\geq1$. This avoids the undefined ecological case of an empty–empty Sørensen comparison.

#### **S1.4 Parameterized richness-matched baselines**

The parameterized baseline does not condition on realized richness. Instead, $y$ is formally changed according to eqn (S10) until expected local richness equals the specified value $\alpha$.

##### **S1.4.1 Coverage deficit**

Under Fisher population weighting, the expected abundance mass contributed by locally absent species is

$$D_{\mathrm{pop}}\left( y \right)=\frac{E_{\phi}\left[ N y^{-N} \right]}{E_{\phi}\left[ N \right]}. \left( S18 \right)$$

Using the generating function in eqn (S2),

$$E_{\phi}\left[ Nt^{N} \right]=t\Psi'\left( t \right),$$

so

$$D_{\mathrm{pop}}\left( y \right)=\frac{\left( x/y \right)/\left( 1-x/y \right)}{x/\left( 1-x \right)}. \left( S19 \right)$$

Substituting the richness bridge in eqn (S10) gives

$$D_{\mathrm{pop}}\left( \alpha\right)=1-\frac{1-\left( 1-x \right)^{\alpha/S}}{x}. \left( S20 \right)$$

Multiplying by the common display factor gives main-text eqn (4),

$$\boxed{B_{\mathrm{CD}}^{\mathrm{par}}\left( \alpha\right)=\frac{M}{M-1}\left[ 1-\frac{1-\left( 1-x \right)^{\alpha/S}}{x} \right].} \left( S21 \right)$$

Equation (S21) uses Fisher population weighting, i.e. the ratio $E\left[ Na\left( N \right) \right]/E\left[ N \right]$. For a finite regional pool whose species abundances are independently drawn from the Fisher distribution, the exact expectation of the normalized quantity $\frac{\sum_{i} N_{i}y^{-N_{i}}}{\sum_{i} N_{i}}$ is not generally equal to this ratio of expectations because the denominator is random. The exact finite-$S$ marginal expectation is

$$D_{\mathrm{fin}}\left( y \right)=S\int_{0}^{1} \frac{1}{y}\Psi'\left( t/y \right)\Psi\left( t \right)^{S-1} dt. \left( S22 \right)$$

Equation (S22) converges to eqn (S19) as the regional pool becomes sufficiently large. The main text retains eqn (S21) because it provides the closed-form population-weighted richness–uniqueness relationship; all conditional baselines below retain the finite-pool normalization exactly.

##### **S1.4.2 Sørensen dissimilarity**

For a randomly selected species, define the first and second occurrence moments

$$\bar{q}=E_{\phi}\left[ q\left( N \right) \right]$$

and

$$\overline{q^{2}}=E_{\phi}\left[ q\left( N \right)^{2} \right].$$

From eqns (S2) and (S5),

$$\bar{q}=1-\frac{\ln\left( 1-x/y \right)}{\ln\left( 1-x \right)}, \left( S23 \right)$$

and

$$\overline{q^{2}}=1-2\frac{\ln\left( 1-x/y \right)}{\ln\left( 1-x \right)}+\frac{\ln\left( 1-x/y^{2} \right)}{\ln\left( 1-x \right)}. \left( S24 \right)$$

For the two-site occurrence indicators of one species,

$$E\left[ t^{X_{i}+Y_{i}} \right]=1-2\bar{q}\left( 1-t \right)+\overline{q^{2}}\left( 1-t \right)^{2}.$$

Define

$$h\left( t \right)=1-2\bar{q}\left( 1-t \right)+\overline{q^{2}}\left( 1-t \right)^{2}. \left( S25 \right)$$

A species contributes to shared richness with probability $\overline{q^{2}}$. Conditional on that species being shared, it contributes two occurrences to $R_{X}+R_{Y}$, while the remaining $S-1$ species contribute according to $h\left( t \right)$. Using

$$\frac{1}{m}=\int_{0}^{1} t^{m-1}dt,$$

the expected Sørensen similarity is

$$E\left( \frac{2A}{R_{X}+R_{Y}} \right)=2S\overline{q^{2}}\int_{0}^{1} t h\left( t \right)^{S-1}dt. \left( S26 \right)$$

Hence

$$\boxed{B_{\mathrm{Sor}}^{\mathrm{par}}=1-2S\overline{q^{2}}\int_{0}^{1} t h\left( t \right)^{S-1}dt.} \left( S27 \right)$$

Under richness matching,

$$\bar{q}=\frac{\alpha}{S},$$

and $y$ is given by eqn (S10), recovering main-text eqn (5).

If desired, $\overline{q^{2}}$ can be written directly in terms of $\alpha$ and $x$. Let

$$r_{\alpha}=1-\left( 1-x \right)^{1-\alpha/S}=\frac{x}{y}.$$

Then

$$\overline{q^{2}}=\frac{2\alpha}{S}-1+\frac{\ln\left( 1-r_{\alpha}^{2}/x \right)}{\ln\left( 1-x \right)}. \left( S28 \right)$$

When $S$ is large enough that $R_{X}+R_{Y}$ is concentrated around its mean, the simpler ratio-of-expectations approximation is

$$B_{\mathrm{Sor}}^{\mathrm{par}}\approx\frac{M}{M-1}\left( 1-\frac{\overline{q^{2}}}{\bar{q}} \right). \left( S29 \right)$$

This approximation is useful for interpretation, whereas eqn (S27) is the exact finite-$S$ expectation under the stated ensemble.

#### **S1.5 Quenched conditioning on realized richness**

The quenched baseline fixes one realized regional abundance vector

$$\mathbf{n}=\left( n_{1},\ldots,n_{S} \right)$$

and therefore fixes

$$p_{i}=\frac{n_{i}}{\sum_{k} n_{k}}, q_{i}=1-y^{-n_{i}}.$$

Only local occurrence remains stochastic.

##### **S1.5.1 Poisson-binomial richness probabilities**

Because species have different occurrence probabilities $q_{i}$, local richness

$$R=\sum_{i=1}^{S} X_{i}$$

has a Poisson-binomial distribution.

For ecological interpretation, it is useful to write its probability directly as

$$P\left( R=r\mid\mathbf{n},y \right)=\sum_{\begin{matrix} \mathcal{A\subseteq\{}1,\ldots,S\} \\ \left| \mathcal{A} \right|=r \end{matrix}} \prod_{i\mathcal{\in A}} q_{i}\prod_{k\mathcal{\notin A}} \left( 1-q_{k} \right). \left( S30 \right)$$

In numerical calculations these probabilities are obtained much more efficiently by recursion. Let $P_{m}\left( r \right)$ be the richness probability after the first $m$ species. Starting from

$$P_{0}\left( 0 \right)=1,$$

the update is

$$P_{m}\left( r \right)=\left( 1-q_{m} \right)P_{m-1}\left( r \right)+q_{m}P_{m-1}\left( r-1 \right). \left( S31 \right)$$

For species $i$, let $R_{-i}$ denote richness contributed by all other species. Bayes’ rule gives

$$P\left( X_{i}=1\mid R=\alpha,\mathbf{n},y \right)=\frac{q_{i}P\left( R_{-i}=\alpha-1 \right)}{P\left( R=\alpha\right)}. \left( S32 \right)$$

Equation (S32) shows explicitly why richness conditioning changes expected species composition: a species is favored not only by its own occurrence probability $q_{i}$, but also by the probability that the remaining species contribute exactly $\alpha-1$ occurrences.

##### **S1.5.2 Coverage-deficit baseline**

Conditional expected coverage is

$$E\left( C_{X}\mid R=\alpha,\mathbf{n},y \right)=\sum_{i=1}^{S} p_{i}P\left( X_{i}=1\mid R=\alpha,\mathbf{n},y \right).$$

Substituting eqn (S32) gives

$$\boxed{B_{\mathrm{CD}}^{Q}\left( \alpha\mid\mathbf{n},y \right)=\frac{M}{M-1}\left[ 1-\sum_{i=1}^{S} p_{i}\frac{q_{i}P\left( R_{-i}=\alpha-1 \right)}{P\left( R=\alpha\right)} \right],} \left( S33 \right)$$

which is main-text eqn (6).

At the upper boundary, $R=S$ implies that every species is represented, so

$$B_{\mathrm{CD}}^{Q}\left( S\mid\mathbf{n},y \right)=0. \left( S34 \right)$$

##### **S1.5.3 Sørensen baseline**

Now consider a focal site $X$ conditioned on

$$R_{X}=\alpha$$

and an independent comparison site $Y$ sampled from the same fixed regional pool.

For species $i$ to contribute to shared richness, it must be present in both sites. The conditional probability of focal presence is given by eqn (S32), while comparison-site presence has probability $q_{i}$. If species $i$ is shared, let $K_{-i}$ be the number of other species present in $Y$. Then

$$R_{Y}=1+K_{-i}$$

and

$$\frac{1}{\alpha+R_{Y}}=\frac{1}{\alpha+1+K_{-i}}.$$

Therefore,

$$E\left[ \frac{1}{\alpha+R_{Y}} | Y_{i}=1 \right]=\sum_{r=0}^{S-1} \frac{P\left( R_{Y,-i}=r \right)}{\alpha+r+1}. \left( S35 \right)$$

Combining these components yields the fully probability-based expression

$$\boxed{B_{\mathrm{Sor}}^{Q}\left( \alpha\mid\mathbf{n},y \right)=1-2\sum_{i=1}^{S} \frac{q_{i}^{2}P\left( R_{X,-i}=\alpha-1 \right)}{P\left( R_{X}=\alpha\right)}\sum_{r=0}^{S-1} \frac{P\left( R_{Y,-i}=r \right)}{\alpha+r+1}.} \left( S36 \right)$$

Because $X$ and $Y$ sample from the same fixed pool, $R_{X,-i}$ and $R_{Y,-i}$ have the same Poisson-binomial distribution. We retain the separate subscripts in eqn (S36) only to clarify their different roles.

Defining the probability-generating function

$$G_{-i}\left( t \right)=\prod_{k\neq i} \left( 1-q_{k}+q_{k}t \right), \left( S37 \right)$$

the final sum in eqn (S36) can equivalently be written as

$$\sum_{r=0}^{S-1} \frac{P\left( R_{Y,-i}=r \right)}{\alpha+r+1}=\int_{0}^{1} t^{\alpha}G_{-i}\left( t \right) dt. \left( S38 \right)$$

Substituting eqn (S38) into eqn (S36) gives main-text eqn (7).

###### Why the quenched Sørensen baseline can turn upward

The high-richness behavior of the quenched Sørensen baseline follows directly from the asymmetric conditioning. The focal richness is fixed at $\alpha$, whereas the richness distribution of the comparison site remains unchanged.

The limiting case $\alpha=S$ is particularly transparent. The focal site then contains every regional species, so every species present in $Y$ is necessarily shared:

$$A=R_{Y}.$$

Consequently,

$$D_{\mathrm{Sor}}=1-\frac{2R_{Y}}{S+R_{Y}}=\frac{S-R_{Y}}{S+R_{Y}},$$

and therefore

$$B_{\mathrm{Sor}}^{Q}\left( S\mid\mathbf{n},y \right)=\frac{M}{M-1}E\left( \frac{S-R_{Y}}{S+R_{Y}} \right). \left( S39 \right)$$

Except in the degenerate case in which the comparison site also contains every species with probability one, eqn (S39) is positive. Thus, unlike coverage deficit, quenched Sørensen uniqueness need not approach zero as focal richness approaches the regional pool size. A minimum followed by an upturn at high richness is therefore possible under strict neutrality.

Whether this behavior is practically relevant depends on the probability support of the richness distribution. In systems where the expected local richness is much smaller than $S$, very high values of $\alpha$ may have extremely small probability even though their conditional expectations remain mathematically well defined.

#### **S1.6 Fisher-averaged annealed conditioning**

The annealed baseline differs from the quenched baseline by treating regional abundance itself as stochastic. The parameters $x$ and $y$ are fixed, but individual regional abundance realizations are integrated over after conditioning on focal richness.

##### **S1.6.1 Posterior regional abundance after focal presence or absence**

For a species observed in the focal site,

$$P\left( N=n\mid X=1 \right)=\frac{\phi\left( n \right)q\left( n \right)}{\bar{q}}, \left( S40 \right)$$

whereas for a species absent from the focal site,

$$P\left( N=n\mid X=0 \right)=\frac{\phi\left( n \right)y^{-n}}{1-\bar{q}}. \left( S41 \right)$$

Thus focal occurrence contains information about regional abundance: species observed locally are posteriorly enriched for larger regional abundances.

The generating function for regional abundance conditional on focal presence is

$$\begin{matrix} U\left( t \right) & =E\left( t^{N}\mid X=1 \right) \\ & =\frac{\Psi\left( t \right)-\Psi\left( t/y \right)}{1-\Psi\left( 1/y \right)} \\ & =\frac{\ln\left( 1-xt \right)-ln\left( 1-xt/y \right)}{\ln\left( 1-x \right)-ln\left( 1-x/y \right)}. \end{matrix} \left( S42 \right)$$

For a species absent from the focal site,

$$\begin{matrix} V\left( t \right) & =E\left( t^{N}\mid X=0 \right) \\ & =\frac{\Psi\left( t/y \right)}{\Psi\left( 1/y \right)} \\ & =\frac{\ln\left( 1-xt/y \right)}{\ln\left( 1-x/y \right)}. \end{matrix} \left( S43 \right)$$

Both satisfy

$$U\left( 1 \right)=V\left( 1 \right)=1.$$

After Fisher averaging, species are exchangeable and each focal occurrence has marginal probability $\bar{q}$. Hence

$$R_{X}\sim Binomial\left( S,\bar{q} \right). \left( S44 \right)$$

Conditional on $R_{X}=\alpha$, every subset of $\alpha$ species is equally likely. For any particular such subset, the regional abundances of the $\alpha$ present species are independent draws from the posterior distribution represented by $U$, while those of the remaining $S-\alpha$ species are independent draws from the distribution represented by $V$.

This exchangeability is what makes the annealed conditional expectations analytically tractable.

##### **S1.6.2 Annealed coverage deficit**

Let

$$N_{+}=\sum_{i=1}^{\alpha} N_{i}^{+}$$

be the total regional abundance of species present in the focal site, where the $N_{i}^{+}$ have generating function $U$, and let

$$N_{-}=\sum_{i=1}^{S-\alpha} N_{i}^{-}$$

be the total abundance of absent species, with generating function $V$.

Focal-site regional coverage is

$$C_{X}=\frac{N_{+}}{N_{+}+N_{-}}. \left( S45 \right)$$

Using the identity

$$\frac{1}{m}=\int_{0}^{1} t^{m-1}dt$$

and symmetry among the $\alpha$ present species,

$$E\left( C_{X}\mid R_{X}=\alpha\right)=\alpha\int_{0}^{1} U'\left( t \right)U\left( t \right)^{\alpha-1}V\left( t \right)^{S-\alpha} dt. \left( S46 \right)$$

Therefore

$$\boxed{B_{\mathrm{CD}}^{A}\left( \alpha;x,y,S,M \right)=\frac{M}{M-1}\left[ 1-\alpha\int_{0}^{1} U'\left( t \right)U\left( t \right)^{\alpha-1}V\left( t \right)^{S-\alpha} dt \right],} \left( S47 \right)$$

which is main-text eqn (8).

An equivalent form computes the deficit directly:

$$B_{\mathrm{CD}}^{A}\left( \alpha;x,y,S,M \right)=\frac{M}{M-1}\left( S-\alpha\right)\int_{0}^{1} U\left( t \right)^{\alpha}V\left( t \right)^{S-\alpha-1}V^{'\left( t \right)} dt. \left( S48 \right)$$

Equations (S47) and (S48) are algebraically identical because the derivative of

$$U\left( t \right)^{\alpha}V\left( t \right)^{S-\alpha}$$

integrates from 0 to 1. Equation (S48) is often numerically preferable when coverage is close to one because it avoids subtracting two nearly equal quantities.

At the boundaries,

$$B_{\mathrm{CD}}^{A}\left( 0 \right)=\frac{M}{M-1}, B_{\mathrm{CD}}^{A}\left( S \right)=0. \left( S49 \right)$$

##### **S1.6.3 Annealed Sørensen baseline**

The same posterior abundance distributions $U$ and $V$ determine the incidence-based expectation.

Because the probability that a species with regional abundance $n$ is absent from an independent comparison site is $y^{-n}$, evaluating a posterior abundance generating function at $t=1/y$ gives its posterior comparison-site absence probability.

For a species present in the focal site,

$$u\equiv U\left( 1/y \right)=P\left( Y=0\mid X=1 \right), \left( S50 \right)$$

and for a species absent from the focal site,

$$v\equiv V\left( 1/y \right)=P\left( Y=0\mid X=0 \right). \left( S51 \right)$$

Explicitly,

$$u=\frac{\ln\left( 1-x/y \right)-ln\left( 1-x/y^{2} \right)}{\ln\left( 1-x \right)-ln\left( 1-x/y \right)}, \left( S52 \right)$$

and

$$v=\frac{\ln\left( 1-x/y^{2} \right)}{\ln\left( 1-x/y \right)}. \left( S53 \right)$$

Thus a focal-present species occurs in the comparison site with probability $1-u$, whereas a focal-absent species occurs there with probability $1-v$.

Conditional on focal richness $\alpha$, shared richness therefore follows

$$A\sim Binomial\left( \alpha,1-u \right),$$

while the number of species occurring only in the comparison site follows

$$B\sim Binomial\left( S-\alpha,1-v \right),$$

with $A$ and $B$ independent. Comparison-site richness is

$$R_{Y}=A+B.$$

Using again the reciprocal integral identity,

$$\begin{matrix} E\left( \frac{2A}{\alpha+A+B} \right) & =2\alpha\left( 1-u \right)\int_{0}^{1} t^{\alpha}\left[ u+\left( 1-u \right)t \right]^{\alpha-1} \\ & \times\left[ v+\left( 1-v \right)t \right]^{S-\alpha} dt. \end{matrix} \left( S54 \right)$$

Therefore

$$\boxed{\begin{matrix} B_{\mathrm{Sor}}^{A}\left( \alpha;x,y,S,M \right) & =1-2\alpha\left[ 1-U\left( \frac{1}{y} \right) \right] \\ & \times\int_{0}^{1} t^{\alpha}\left[ U\left( \frac{1}{y} \right)+\left\{ 1-U\left( \frac{1}{y} \right) \right\}t \right]^{\alpha-1}\left[ V\left( \frac{1}{y} \right)+\left\{ 1-V\left( \frac{1}{y} \right) \right\}t \right]^{S-\alpha}dt \end{matrix}}, \left( S55 \right)$$

which is main-text eqn (9).

Equations (S47) and (S55) therefore arise from the same two posterior abundance distributions. Coverage deficit uses their complete abundance distributions because regional abundance weights enter the metric directly, whereas Sørensen needs only the posterior probabilities of occurrence in the comparison site, obtained by evaluating $U$ and $V$ at $1/y$.

Because focal presence favors species with larger regional abundances,

$$U\left( 1/y \right)<V\left( 1/y \right)$$

for a nondegenerate Fisher distribution, and hence

$$P\left( Y=1\mid X=1 \right)>P\left( Y=1\mid X=0 \right). \left( S56 \right)$$

This posterior coupling is absent from a fixed-pool interpretation and helps explain why annealed and quenched Sørensen curves can differ substantially.

#### **S1.7 Relationship between quenched and annealed conditioning**

The annealed baseline is not an unweighted average of quenched baselines.

Let

$$B^{Q}\left( \alpha\mid\mathbf{N} \right)$$

denote either quenched conditional uniqueness score for a regional abundance realization $\mathbf{N}$, and let

$$P_{\mathbf{N}}\left( \alpha\right)=P\left( R_{X}=\alpha\mid\mathbf{N} \right)$$

be the probability that this regional state generates focal richness $\alpha$.

Bayes’ rule gives

$$P\left( \mathbf{N}\mid R_{X}=\alpha\right)\propto P_{\mathbf{N}}\left( \alpha\right)P\left( \mathbf{N} \right).$$

Consequently,

$$B^{A}\left( \alpha\right)=\frac{E_{\mathbf{N}}\left[ P_{\mathbf{N}}\left( \alpha\right)B^{Q}\left( \alpha\mid\mathbf{N} \right) \right]}{E_{\mathbf{N}}\left[ P_{\mathbf{N}}\left( \alpha\right) \right]}. \left( S57 \right)$$

Regional abundance configurations that are more likely to produce the observed richness therefore receive more weight in the annealed expectation.

Equation (S57) provides a direct Monte Carlo interpretation of the annealed model. If many independent Fisher regional pools are generated and a quenched conditional curve is calculated for each, their simple arithmetic mean does not estimate the annealed baseline. Instead, at each richness $\alpha$, each quenched curve must be weighted by the probability that its regional pool generates that richness.

This identity also clarifies the difference between the three baselines:

- the parameterized baseline changes the occurrence regime until $E\left( R \right)=\alpha$;
- the quenched baseline fixes $\left( \mathbf{n},y \right)$ and conditions on $R=\alpha$;
- the annealed baseline fixes $\left( x,y \right)$, conditions on $R=\alpha$, and then integrates over regional abundance realizations with the posterior weights in eqn (S57).

They are therefore different expectations, not alternative computational approximations to the same quantity.
