## Appendix S2 for "A sampling-theory framework for richness–uniqueness relationships"

Appendix S2: Supplementary figures and tables

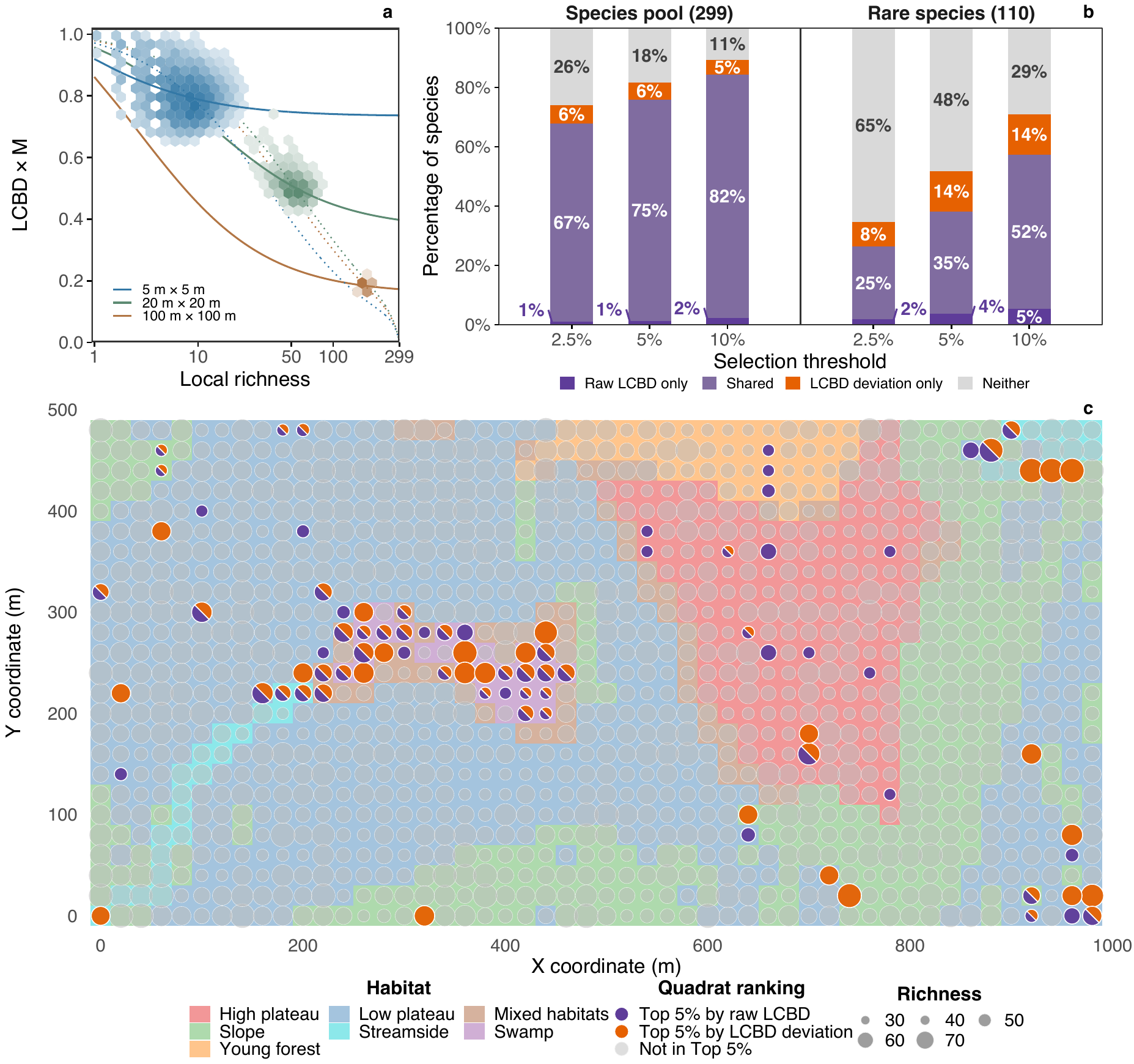

Figure S1. Richness–uniqueness relationships in the BCI forest plot. (a) Sørensen LCBD versus local richness across three spatial grains. Hexagon shading indicates quadrat density; dotted and solid curves show fitted parameterized and annealed conditional expectations, respectively. (b) Representation of the regional species pool and rare species by 20 m ×20 m quadrats selected under raw-LCBD or LCBD-deviation rankings at alternative thresholds. (c) Spatial distribution of quadrats in the top 5% of the two rankings at the 20 m × 20 m grain.

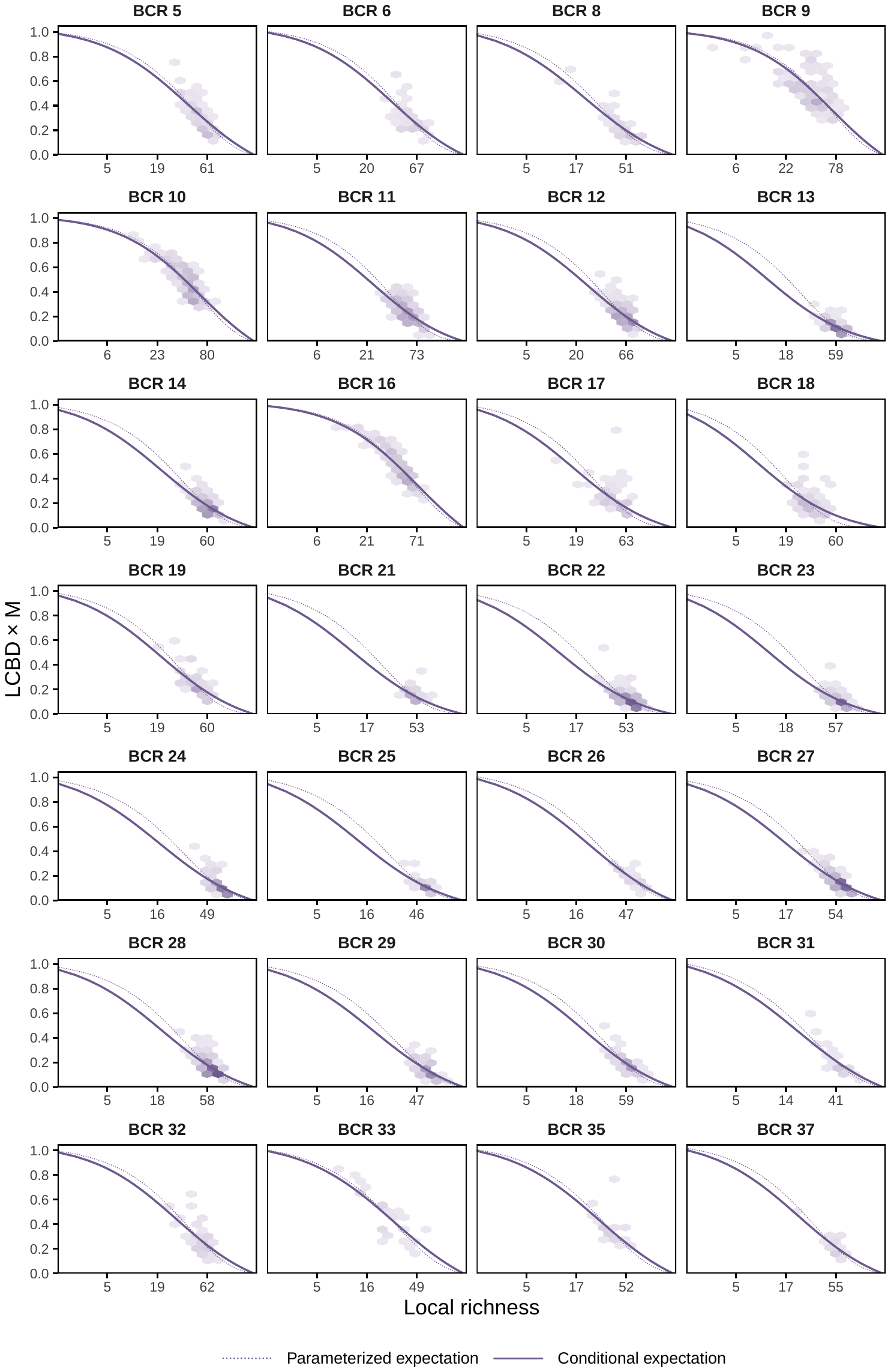

**Figure S2 Coverage-deficit uniqueness versus richness across North American BBS Bird Conservation Regions.** Hexagon shading indicates the route density.

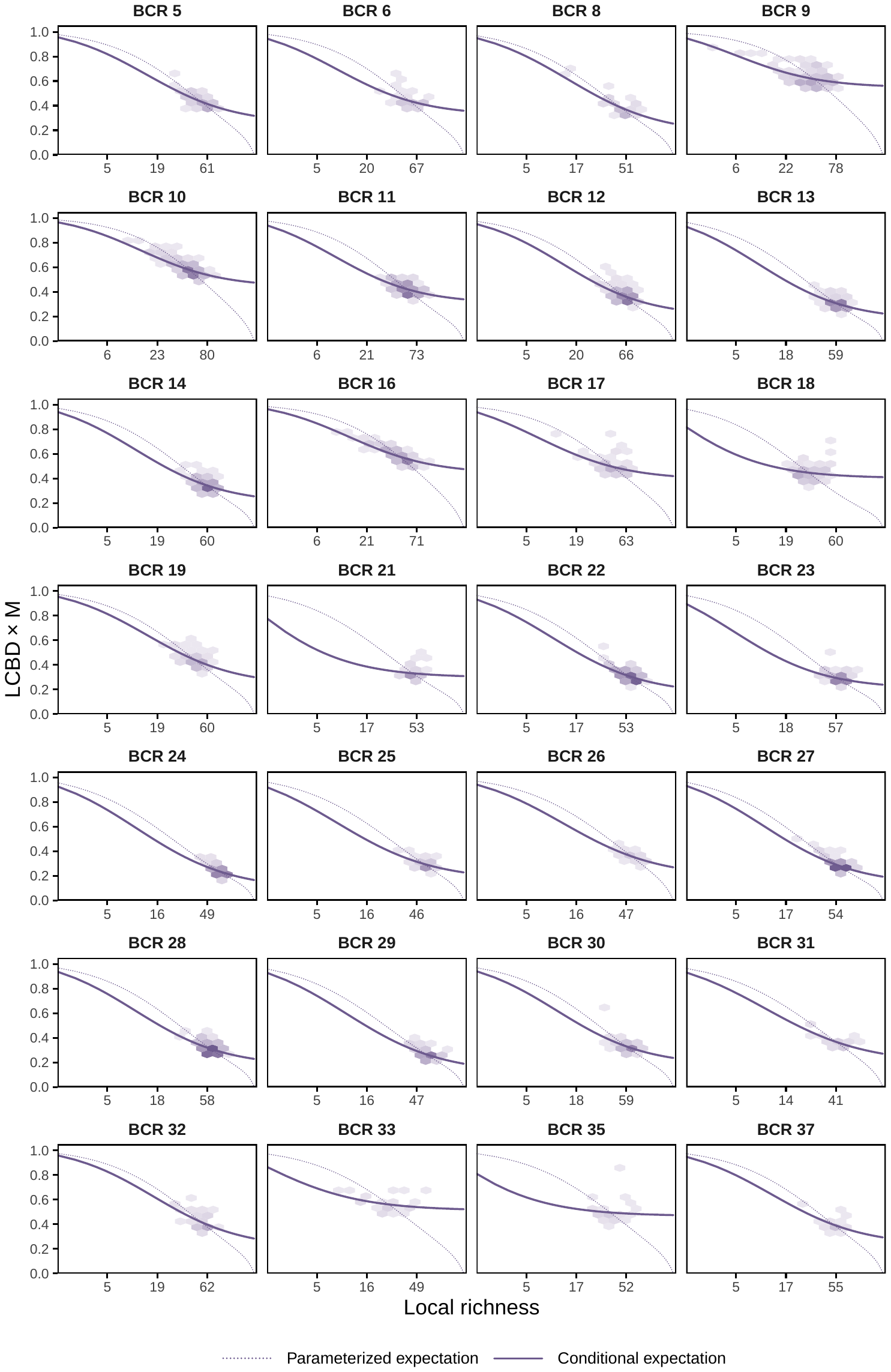

**Figure S3 Sørensen uniqueness versus richness across North American BBS Bird Conservation Regions.** Hexagon shading indicates the route density.

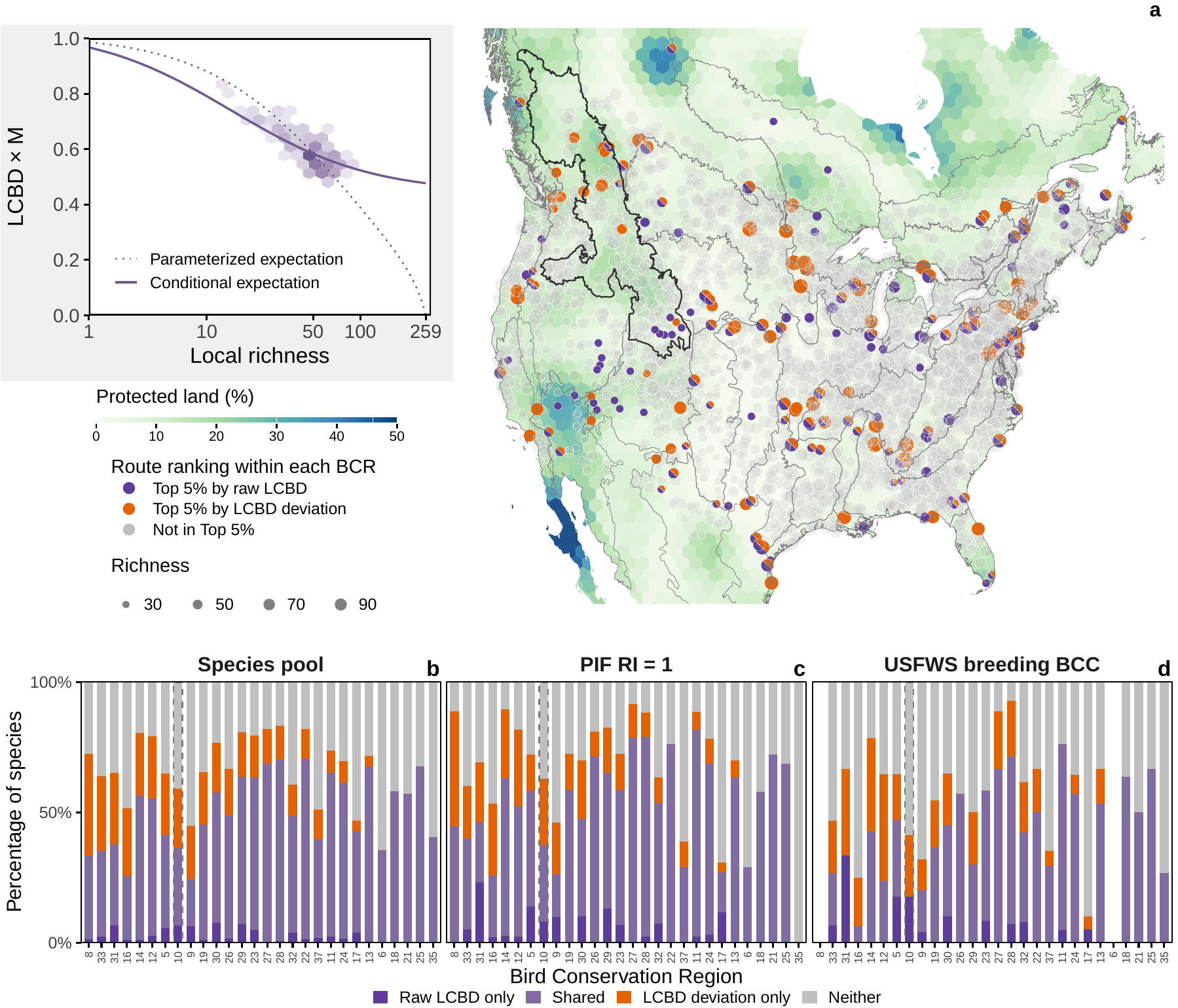

Figure S4. Richness–uniqueness relationships across North American BBS Bird Conservation Regions. (a) BBS routes in the top 5% of raw-LCBD and LCBD-deviation rankings within each BCR, shown over protected-land coverage on a 100-km-spaced hexagonal grid. The inset shows Sørensen uniqueness versus richness in the Northern Rockies (BCR 10, highlighted across panels). (b–d) Representation of the regional species pool (b), PIF regionally important species (c), and USFWS breeding Birds of Conservation Concern (d) under the two rankings.

**Table S1. Representation of the regional species pool and regionally listed species among the top 5% of BBS routes within North American BBS Bird Conservation Region under the coverage-deficit framework.**

| **BCR code** | **Bird Conservation Region** | **Number of species** | | | **Species represented by top 5% of routes (%)** | | | | | |
| --- | --- | --- | --- | --- | --- | --- | --- | --- | --- | --- |
|  |  | **Regional species pool** | **PIF**  **RI = 1** | **USFWS**  **BCC** | **Raw LCBD** | | | **LCBD deviation** | | |
|  |  |  |  |  | **Regional species pool** | **PIF**  **RI = 1** | **USFWS**  **BCC** | **Regional species pool** | **PIF**  **RI = 1** | **USFWS BCC** |
| 5 | Northern Pacific Rainforest | 182 | 36 | 17 | 44.5 | 55.6 | 58.8 | 45.1 | 52.8 | 52.9 |
| 6 | Boreal Plains | 203 | 31 | — | 32.0 | 29.0 | — | 32.0 | 29.0 | — |
| 8 | Boreal Softwood Shield | 145 | 27 | — | 33.1 | 44.4 | — | 39.3 | 48.1 | — |
| 9 | Great Basin | 250 | 61 | 25 | 26.0 | 27.9 | 24.0 | 38.0 | 36.1 | 24.0 |
| 10 | Northern Rockies | 259 | 62 | 17 | 36.3 | 37.1 | 17.6 | 55.2 | 51.6 | 35.3 |
| 11 | Prairie Potholes | 227 | 44 | 21 | 61.2 | 77.3 | 66.7 | 69.6 | 84.1 | 76.2 |
| 12 | Boreal Hardwood Transition | 201 | 44 | 17 | 43.3 | 38.6 | 23.5 | 75.6 | 72.7 | 58.8 |
| 13 | Lower Great Lakes/St. Lawrence Plain | 173 | 30 | 15 | 64.2 | 60.0 | 46.7 | 68.2 | 66.7 | 53.3 |
| 14 | Atlantic Northern Forest | 179 | 38 | 14 | 64.8 | 68.4 | 57.1 | 76.5 | 81.6 | 85.7 |
| 16 | Southern Rockies/Colorado Plateau | 221 | 47 | 16 | 19.9 | 25.5 | 6.2 | 35.7 | 31.9 | 25.0 |
| 17 | Badlands and Prairies | 190 | 26 | 20 | 45.8 | 30.8 | 15.0 | 43.2 | 19.2 | 5.0 |
| 18 | Shortgrass Prairie | 177 | 19 | 10 | 44.1 | 52.6 | 46.4 | 57.6 | 57.9 | 54.5 |
| 19 | Central Mixed Grass Prairie | 179 | 29 | 12 | 36.9 | 37.9 | 18.2 | 39.1 | 41.4 | 27.3 |
| 21 | Oaks and Prairies | 149 | 18 | 6 | 51.7 | 61.1 | 16.7 | 55.0 | 66.7 | 33.3 |
| 22 | Eastern Tallgrass Prairie | 150 | 21 | 12 | 75.3 | 71.4 | 58.3 | 76.7 | 71.4 | 58.3 |
| 23 | Prairie Hardwood Transition | 166 | 29 | 12 | 57.8 | 48.3 | 50.0 | 65.1 | 72.4 | 75.0 |
| 24 | Central Hardwoods | 135 | 32 | 14 | 65.9 | 75.0 | 64.3 | 61.5 | 75.0 | 57.1 |
| 25 | West Gulf Coastal Plain/Ouachitas | 127 | 35 | 11 | 49.6 | 62.9 | 58.3 | 67.7 | 68.6 | 63.6 |
| 26 | Mississippi Alluvial Valley | 129 | 21 | 7 | 51.2 | 71.4 | 57.1 | 57.4 | 85.7 | 71.4 |
| 27 | Southeastern Coastal Plain | 156 | 47 | 18 | 60.9 | 66.0 | 55.6 | 69.9 | 80.9 | 66.7 |
| 28 | Appalachian Mountains | 168 | 43 | 14 | 66.1 | 79.1 | 71.4 | 72.0 | 79.1 | 78.6 |
| 29 | Piedmont | 129 | 23 | 10 | 68.2 | 69.6 | 60.0 | 67.4 | 73.9 | 80.0 |
| 30 | New England/Mid-Atlantic Coast | 172 | 40 | 20 | 58.1 | 52.5 | 55.0 | 58.1 | 52.5 | 55.0 |
| 31 | Peninsular Florida | 109 | 13 | 9 | 37.6 | 46.2 | 33.3 | 37.6 | 46.2 | 33.3 |
| 32 | Coastal California | 185 | 41 | 26 | 37.3 | 41.5 | 38.5 | 42.2 | 46.3 | 34.6 |
| 33 | Sonoran and Mojave Deserts | 136 | 20 | 14 | 11.0 | 10.0 | 7.1 | 47.8 | 50.0 | 21.4 |
| 35 | Chihuahuan Desert | 148 | 15 | 15 | 40.5 | 0.0 | 26.7 | 52.0 | 26.7 | 60.0 |
| 37 | Gulf Coastal Prairie | 159 | 31 | 17 | 39.6 | 29.0 | 29.4 | 49.1 | 38.7 | 35.3 |

An em dash indicates that USFWS breeding BCC classification was unavailable.

**Table S2. Representation of the regional species pool and regionally listed species among the top 5% of BBS routes within North American BBS Bird Conservation Region under the Sørensen framework.**

| **BCR code** | **Bird Conservation Region** | **Number of species** | | | **Species represented by top 5% of routes (%)** | | | | | |
| --- | --- | --- | --- | --- | --- | --- | --- | --- | --- | --- |
|  |  | **Regional species pool** | **PIF**  **RI = 1** | **USFWS BCC** | **Raw LCBD** | | | **LCBD deviation** | | |
|  |  |  |  |  | **Regional species pool** | **PIF**  **RI = 1** | **USFWS BCC** | **Regional species pool** | **PIF**  **RI = 1** | **USFWS BCC** |
| 5 | Northern Pacific Rainforest | 182 | 36 | 17 | 41.2 | 58.3 | 47.1 | 59.3 | 58.3 | 47.1 |
| 6 | Boreal Plains | 203 | 31 | — | 35.5 | 29.0 | — | 35.5 | 29.0 | — |
| 8 | Boreal Softwood Shield | 145 | 27 | — | 33.1 | 44.4 | — | 71.0 | 88.9 | — |
| 9 | Great Basin | 250 | 61 | 25 | 24.4 | 26.2 | 20.0 | 38.4 | 36.1 | 28.0 |
| 10 | Northern Rockies | 259 | 62 | 17 | 36.3 | 37.1 | 17.6 | 52.5 | 54.8 | 23.5 |
| 11 | Prairie Potholes | 227 | 44 | 21 | 65.2 | 81.8 | 76.2 | 71.4 | 86.4 | 71.4 |
| 12 | Boreal Hardwood Transition | 201 | 44 | 17 | 55.2 | 52.3 | 23.5 | 76.6 | 79.5 | 64.7 |
| 13 | Lower Great Lakes/St. Lawrence Plain | 173 | 30 | 15 | 67.6 | 63.3 | 53.3 | 71.1 | 70.0 | 66.7 |
| 14 | Atlantic Northern Forest | 179 | 38 | 14 | 56.4 | 63.2 | 42.9 | 79.3 | 86.8 | 78.6 |
| 16 | Southern Rockies/Colorado Plateau | 221 | 47 | 16 | 25.3 | 25.5 | 6.2 | 50.7 | 51.1 | 25.0 |
| 17 | Badlands and Prairies | 190 | 26 | 20 | 42.6 | 26.9 | 5.0 | 43.2 | 19.2 | 5.0 |
| 18 | Shortgrass Prairie | 177 | 19 | 10 | 58.2 | 57.9 | 63.6 | 58.2 | 57.9 | 63.6 |
| 19 | Central Mixed Grass Prairie | 179 | 29 | 12 | 45.3 | 58.6 | 36.4 | 64.2 | 72.4 | 54.5 |
| 21 | Oaks and Prairies | 149 | 18 | 6 | 57.0 | 72.2 | 50.0 | 57.0 | 72.2 | 50.0 |
| 22 | Eastern Tallgrass Prairie | 150 | 21 | 12 | 70.7 | 76.2 | 50.0 | 80.7 | 76.2 | 66.7 |
| 23 | Prairie Hardwood Transition | 166 | 29 | 12 | 63.3 | 58.6 | 58.3 | 74.7 | 65.5 | 50.0 |
| 24 | Central Hardwoods | 135 | 32 | 14 | 61.5 | 68.8 | 57.1 | 68.1 | 75.0 | 64.3 |
| 25 | West Gulf Coastal Plain/Ouachitas | 127 | 35 | 11 | 67.7 | 68.6 | 66.7 | 67.7 | 68.6 | 66.7 |
| 26 | Mississippi Alluvial Valley | 129 | 21 | 7 | 48.8 | 71.4 | 57.1 | 65.1 | 81.0 | 57.1 |
| 27 | Southeastern Coastal Plain | 156 | 47 | 18 | 68.6 | 78.7 | 64.7 | 82.1 | 91.5 | 88.9 |
| 28 | Appalachian Mountains | 168 | 43 | 14 | 70.2 | 79.1 | 71.4 | 82.7 | 86.0 | 85.7 |
| 29 | Piedmont | 129 | 23 | 10 | 63.6 | 65.2 | 30.0 | 73.6 | 69.6 | 55.6 |
| 30 | New England/Mid-Atlantic Coast | 172 | 40 | 20 | 57.6 | 47.5 | 45.0 | 69.2 | 60.0 | 55.0 |
| 31 | Peninsular Florida | 109 | 13 | 9 | 37.6 | 46.2 | 33.3 | 58.7 | 46.2 | 33.3 |
| 32 | Coastal California | 185 | 41 | 26 | 48.6 | 53.7 | 42.3 | 56.8 | 56.1 | 50.0 |
| 33 | Sonoran and Mojave Deserts | 136 | 20 | 14 | 35.3 | 40.0 | 28.6 | 61.8 | 55.0 | 42.9 |
| 35 | Chihuahuan Desert | 148 | 15 | 15 | 40.5 | 0.0 | 26.7 | 40.5 | 0.0 | 26.7 |
| 37 | Gulf Coastal Prairie | 159 | 31 | 17 | 39.6 | 29.0 | 29.4 | 49.1 | 38.7 | 35.3 |

An em dash indicates that USFWS breeding BCC classification was unavailable.
